# Linalool fumigation improves mating competitiveness of males for population suppression of the global fruit pest *Cydia pomonella*

**DOI:** 10.1101/2024.07.21.604520

**Authors:** Sheng-Wang Huang, Peng-Cheng Wang, Yan Wang, Jie-Qiong Wang, Ping Gao, Qing-E Ji, Xue-Qing Yang

**Author notes:** Correspondence (P.G.) (X.Q.Y). These authors contributed equally to this work (Sheng-Wang Huang, Peng-Cheng Wang).

## Abstract

**BACKGROUND:** The implementation of sterile insect technique (SIT) has proven effective in the area-wide suppression of several significant agricultural and sanitary pests by employing traditional cobalt-60 (^60^ Co-γ) as a radiation source. Recently, X-ray has been validated as a feasible alternative to ^60^ Co-γ radiation sources. Nonetheless, higher doses of X-ray irradiation lead to insect sterility but diminish mating competitiveness, thereby impacting the effectiveness of SIT applications. Thus, it is crucial to ascertain the optimal irradiation dose and develop strategies to enhance the mating competitiveness of sterile insects to enhance SIT efficacy.

**RESULTS:** In this study, we determined the effect of various X-ray irradiation doses (ranging from 0 to 366 Gy) on the fecundity, fertility, and mating competitiveness of *Cydia pomonella*, a globally invasive fruit pest. Results demonstrated that the sterility rate of sterile males increased proportionally with irradiation dose up to 200 Gy, beyond which it plateaued. Notably, exposure to 200 Gy of irradiation notably decreased the mating competitiveness of male, as evidenced by a mating competitiveness index of 0.17 in laboratory and 0.096 in the orchard. This decline in mating competitiveness is likely linked to the down-regulation of genes associated with the recognition of sex pheromones, specifically *CpomOR3a*, *CpomOR3b*, and *CpomOR5*, following X-ray irradiation. Fumigation of the plant volatile, linalool at varying concentrations (70, 83, and 96 μ L/m ³) resulted in differential enhancements in male mating competitiveness, with the moderate concentration significantly improving the competitiveness of sterilized males, possibly by restoring their ability to recognize sex pheromones. Implementation of repeated releases of sterilized males on a pilot scale led to a notable reduction in the population of *C. pomonella* in the field.

**CONCLUSION:** These findings indicate that fumigation with plant volatiles has the potential to mitigate male sterility induced by X-ray irradiation, offering a promising approach to enhance the efficacy of SIT applications for the control of *C. pomonella*.

**Graphic Abstract:** 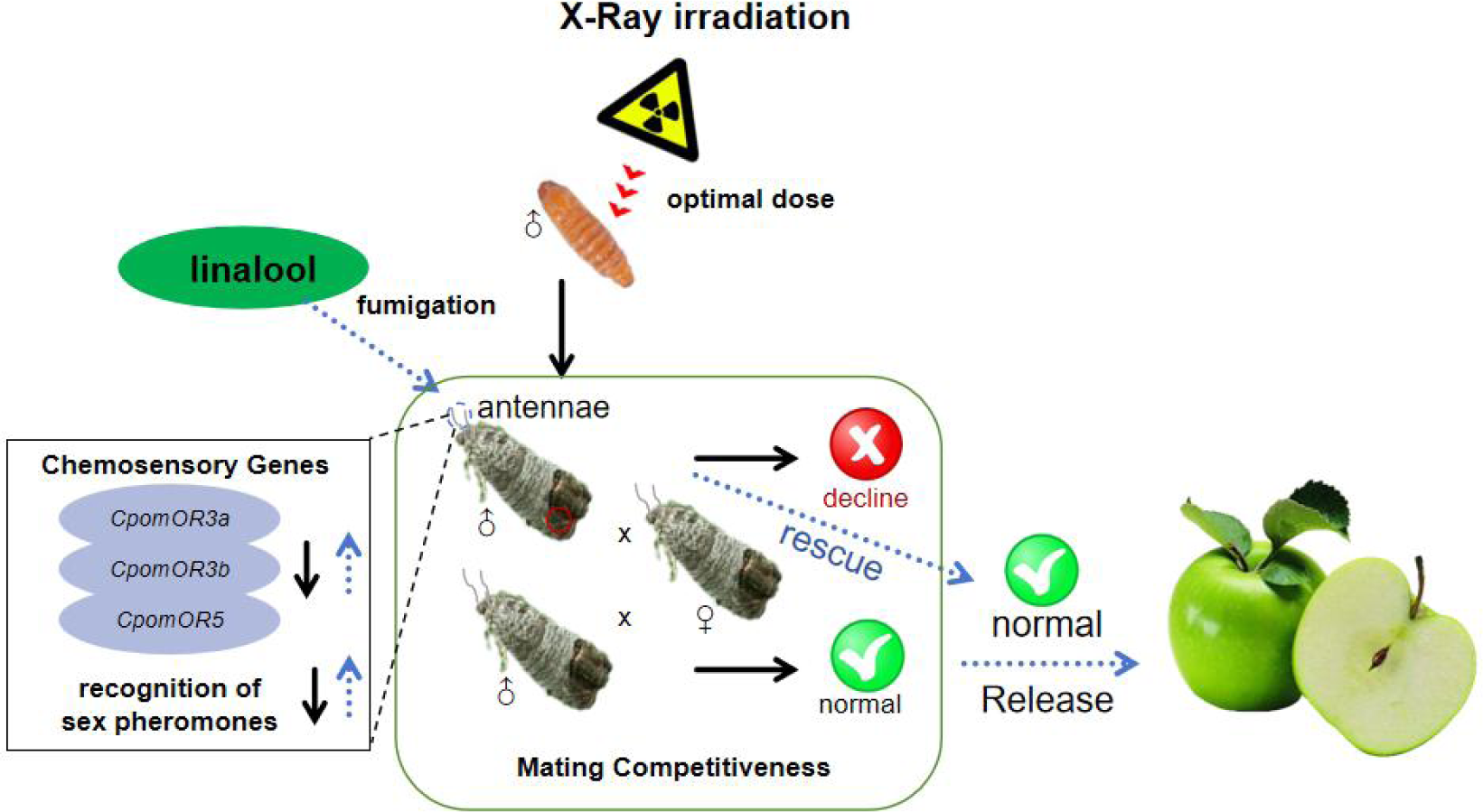

We determined the optimal X-ray irradiation dose and developed a linalool fumigation approach to improve the mating competitiveness of sterilized insects, thereby bolstering the efficacy of against *Cydia pomonella*.

## 1. Introduction

The codling moth, *Cydia pomonella* (Linnaeus), is a notorious pest that infests pome fruits and walnuts worldwide^1^. Various methods, including agricultural, mechanical, chemical, and biological controls, have been employed to manage the population of *C. pomonella*.^2^ Despite efforts to utilize sustainable practices such as agricultural techniques, pheromone-based mating disruption, and attract-kill strategies, effectively controlling dense populations of this pest remains a challenge.^2^ Traditionally, the predominant approach to combat *C. pomonella* has been through the use of conventional insecticides^3^. However, the efficacy of chemical insecticides in controlling this pest is limited due to factors like overlapping generations and the larvae’s habit of tunneling into fruit.^4,5^ Furthermore, the widespread application of insecticides has led to the development of resistance in *C. pomonella*.^5^

Insect sterility technique (SIT) is recognized as an eco-friendly, efficient, and sustainable approach incorporated within the broader framework of Integrated Pest Management (IPM).^6^ The utilization of irradiated males in conjunction with non-irradiated females results in sterile offspring.^7^ However, the slower progress in developing sterile insect control methods for Lepidoptera in comparison to Diptera is linked to the former’s resistance to radiation.^8^ This resistance in Lepidopteran insects is likely due to their necessity for higher radiation doses for sterilization, which can compromise their mating competitiveness and impede the success of sterile insect initiatives. Nevertheless, the effective implementation of the Okanagan- Kootenay Sterile Insect Release (OK SIR) program in British Columbia, Canada led to the successful eradication of *C. pomonella* populations by strategically employing conventional cobalt-60 (^60^ Co-γ) source in area-wide pest management activities.^9^ Ongoing endeavors in insect SIT focus on assessing the effects of radiation on insect fertility, longevity, flight capacity, and mating competitiveness,^10^ with the aim of establishing standardized procedures for consistent and reproducible induction of sterility.^11^

The performance of irradiated insects, particularly their mating competitiveness and flight capacity, is of significant importance in the SIT program, as it directly impacts the dispersal range and mating effectiveness of sterile insects in natural environments.^12^ Previous research has indicated that male pupae irradiated with X-rays at a dose of 366 Gy resulted in complete sterility in *C. pomonella*, albeit with a low mating competitiveness index of 0.01.^13^ The observed side effects of X-ray irradiation, such as reduced mating competitiveness, are likely attributed to its negative impact on flight ability.^4^ Consequently, it is imperative to optimize irradiation doses and devise strategies to improve both mating competitiveness and flight capacity of sterile insects to ensure the effective implementation of the SIT program for pest management.

Host plant volatiles have been documented as influential factors in various behaviors exhibited by phytophagous insects, encompassing mating, feeding, fertilization, directed flight, and pheromone production.^14^ Linalool, a volatile compound categorized as a light-flavored terpene alcohol,^15^ has exhibited a significant attraction towards adult *C. pomonella*, leading to a considerable electrophysiological reaction from their antennae.^16^ Studies conducted on *Ceratitis capitata* have shown an improvement in mating competitiveness following linalool fumigation, while not impacting the survival rate of sterile males.^15^ Nevertheless, the influence of linalool fumigation on the mating competitiveness of male *C. pomonella* remains unexplored.

The olfactory system plays a vital role in various insect behaviors like feeding, orientation, host-seeking, mating, and oviposition.^17,18^

Chemoreception in insects is a complex process involving various chemosensory genes, such as odorant receptors (ORs), gustatory receptors (GRs), ionotropic receptors (IRs), odorant-binding proteins (OBPs), chemosensory proteins (CSPs), and sensory neuron membrane proteins (SNMPs).^19^ In particular, the importance of chemosensory genes in recognizing pheromones in *C. pomonella* has been extensively researched. Tian *et al*.^20^ have identified male pheromone receptor genes in *C. pomonella*, including *CpomOR1*, *CpomOR2a*, *CpomOR3*, *CpomOR5*, *CpomOR6a*, and *CpomOR7*, through transcriptome analysis of the antenna. Knock-out the *CpomOR1* gene using CRISPR/Cas9 technology has been shown to impact the fertility and fecundity of *C. pomonella*.^21^ Moreover, *CpomOR3a* and *CpomOR3b* genes in *C. pomonella* are involved in recognizing plant volatiles and the sex pheromone codlemone, thereby enhancing the moths’ response to codlemone.^22^ However, the underlying cause of the diminished mating competitiveness observed in irradiated *C. pomonella* is yet to be determined, specifically regarding its potential interference with the expression of sex pheromone-related genes. Furthermore, the efficacy of linalool fumigation in mitigating these effects remains to be elucidated.

The objective of this research is to: 1) ascertain the most effective irradiation dosage to achieve a harmonious equilibrium between sterility and mating competitiveness; 2) develop strategies to enhance the mating competitiveness of sterile insects; 3) elucidate the potential mechanism of the decline in mating competitiveness caused by irradiation and improvement following linalool fumigation; 4) investigate the effect of pilot-release of sterile males on suppression of the *C. pomonella* field population.

## 2. Materials and methods

### 2.1 Insects

The *C. pomonella* strain utilized in the investigation was initially obtained from the Laboratory of Agricultural Invasive Biological Prevention and Monitoring at the Institute of Plant Protection, Chinese Academy of Agricultural Sciences. This strain was reared for more than 70 generations within a growth chamber (MLR-352H-PC; Panasonic, Osaka, Japan) under controlled conditions devoid of irradiation. The rearing was maintained a temperature of 26 ± 1 ° C, relative humidity ranging from 60% ± 5%, and a photoperiod set at 16 hours of light followed by 8 hours of darkness.^23^

### 2.2 Irradiation

An X-ray irradiator (SR 100, Changzhou Sairui Instrument Technology Co., Ltd, Changzhou, China) that was furnished with a Radcal Accu-Dose+ digitizer featuring a 10 × 6–0.6 CT ion chamber for the purpose of measuring the average dose rate, was utilized in this study. This instrument was employed to expose 8-day-old male pupae (1 day prior to emergence),^13^ which were positioned within a cylindrical transparent tube (diameter: 2 cm; length: 6 cm) located in the irradiation chamber (dimensions: length: 21 cm; width: 24 cm; height: 16 – 40 cm) to X-ray radiation at a dose rate of 4.3 Gy/min.

### 2.3 Effects of different doses of X-ray irradiation on the mating and reproduction of *C. pomonella*

The 8-day-old male pupae were divided into five groups comprising 90 individuals each, and subjected to X-ray irradiation at doses of 0, 50, 100, 150, 200, 250, and 300 Gy, respectively. Subsequently, the pupae within each treatment were randomly divided into three groups consisting of 30 pupae each, and placed in an incubator for emergency situations, with the subsequent emergence count being documented. The matured males and females from the same day were introduced into a mating box (25*12*15 cm) at a ratio of 10 females to 10 males, and each treatment was repeated seven times. Following the laying of eggs by the females and subsequent collection of the egg papers, they were placed in individual disposable transparent plastic bags (15*26 cm) alongside damp wipes on a daily basis, and then transferred to the incubator for hatching. The eggs laid was recorded and the hatching rate was calculated upon the demise of all females. The mortality of the adult was monitored per day, with deceased individuals promptly removed. The lifespan of males was documented, and expired females were gathered in centrifuge tubes (1.5 mL) for dissection under a dissecting microscope to observe mating sac morphology, define mating status, and calculate female mating rates according to Zhang et al.^13^

Different proportions of “irradiated male: non-irradiated male: non-irradiated female” at ratios of 0:10:10, 10:0:10, and 10:10:10 were established, respectively, to calculate the mating competition index. These male individuals, both irradiated and non-irradiated, along with the female counterparts, were matched in mating age(25*12*15 cm). The eggs laid (fecundity) and hatched (fertility) were meticulously documented for each grouping, with every treatment being replicated thrice for statistical robustness. The competitive mating index (C) of sterile males was then determined utilizing the methodology outlined by Fried.^24^

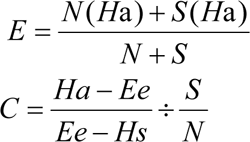

*N* is the number of normal male moths and *S* is the number of irradiated male moths. *Ha* is the hatching rate of normal male moths paired with normal female moths. *Hs* is the hatching rate of sterile male moths paired with normal female moths. *Ee* is the hatching rate of mixed male moths of a certain S/N ratio paired with normal female moths. *E* is the expected hatching rate of a mixed S/N ratio male moth paired with a normal female moth.

### 2.4 Effect of linalool fumigation on mating competitiveness of *C. pomonella*

Three disposable transparent plastic boxes (21*14*7 cm) with a volume of 1500 mL each were assembled for experimentation. A strip of filter paper (1 cm in wide and 5 cm long) was suspended in the center of the lid of each box. Male moths that newly emerged were introduced into the boxes.

Subsequently, linalool was evenly distributed onto the filter paper strips within the boxes to achieve concentrations of 70, 83, and 96 μ L/m ³, which were clearly labeled on the lids of the respective boxes. After a 3-hour fumigation period, the moths were removed and the males and females from the same day were segregated into adult mating boxes at a 1:1 ratio. The assessment procedures remained consistent with those described in section 2.3.

### 2.5 Effect of linalool fumigation on mating competitiveness of sterile *C. pomonella*

To investigate the effect of linalool fumigation on mating competitiveness of sterile *C. pomonella* in laboratory, the 8-day-old male pupae were irradiated with X-rays at a dose of 200 Gy, subsequently maintained in a growth chamber until emergence. The freshly emerged male adults underwent a 3- hour fumigation with linalool. The procedures for fumigation, assessment of average survival time, fecundity per female, egg hatchability, mating frequency, and mating competitiveness followed the protocols described above- mentioned.

To investigate the effect of linalool fumigation on mating competitiveness of sterile *C. pomonella* in orchard, a field trial was conducted at Wumeifang Village, Jiuzhai Town, Gaizhou City, Yingkou City, Liaoning Province (40°3′52 ”N, 122 ° 5 ′ 44 ”E) from August 1 to September 2, 2023. Small insect rearing nets (60 cm*60 cm*90 cm) with 40 mesh were affixed to tree trunks, and adult mating age were placed inside these nets for fumigation purposes. The experimental procedures for evaluating mean longevity, single female fecundity, hatching rate, mating rate, and mating competitiveness were consistent with those described above.

### 2.6 Effects of linalool fumigation on the flight ability of sterile *C. pomonella* moths

The flight performance of moths was assessed using the FXMD-24-USB insect flight information system (Henan Hebi Jiaduoke Industry and Trade Co., Ltd, Hebi, Henan, China), according to the method described in Huang et al.^4^ Normal, sterile, and fumigated sterile male moths were segregated into distinct adult rearing boxes and provided with a daily diet of 10% honey water. Then, 25 healthy male moths with intact wings that were 3 days post- emergence were selected for the study, and the flight capability of each group was evaluated. The insects were initially immobilized on a chilled surface before their abdomens were exposed. A copper wire was fashioned into a loop, folded at a 90° angle, and affixed to the moth’s abdomen using a small amount of adhesive, ensuring proper fixation by inflation and rapid solidification of the glue. The other end of the wire was secured to the arm of the flight mill, positioning the moth parallel to the ground and aligning its flight direction with the mill. Subsequently, the insect flight information system was activated to input data on the test subjects. Flight ability assessments were conducted from 22:00 to 6:00 in a darkened environment, maintaining a constant ambient temperature of 25 ± 1 ℃ and relative humidity of 70 ± 5% throughout the evaluation process.

### 2.7 RNA isolation and cDNA synthesis

Total RNA was isolated from a collective of ten 3-day-old male adults of *C. pomonell*a, who were subjected to either 200 Gy X-ray irradiation or not, utilizing the EAStep Super Total RNA Extraction Kit (Promega, Shanghai, China) in accordance with the manufacturer’s guidelines. The concentration and quality of the RNA samples were assessed employing a NanoDrop2000 spectrophotometer (Thermo Fisher Scientific, Waltham, MA, USA).

Subsequently, 1 μg of RNA from each treatment was utilized in the synthesis of complementary DNA (cDNA) using the GoScript ™ Reverse Transcription Mix (Promega, Madison, WI, USA). The resulting cDNA was preserved at - 20 °C until further analysis.

### 2.8 Reverse-transcription quantitative polymerase chain reaction analysis

The mRNA levels of pheromone recognition-related genes (Table S1, Figure S1-S8) were measured using reverse transcription quantitative polymerase chain reaction (RT-qPCR) with Bio-Rad CFX96 (Bio-Rad, Hercules, CA, USA) in a reaction of 20 μ L, containing of 1 μ L of cDNA template, 10 μ L of TB Green Premix Ex Taq 2 (TliRNaseH Plus) (TaKaRa, Dalian, China), 1 μ L of each primer (10 μ mol/L), and 7 μ L double-distilled water (ddH2O). The RT-qPCR cycled at 95 °C for 30 s, followed by 40 cycles of 95 °C for 5 s and 60 °C for 30 s. Afterward, a melting curve was conducted starting at 55– 95 ° C to identify the specificity of PCR products. The internal control genes used were *EF-1α* (MN037793) and *β-Actin* (KC832921) of *C. pomonella*.^25^ For each three biological replicates and three technical replicates were conducted. Relative expression levels for each gene were calculated using the 2^−△△CT^ method.^26^

### 2.9 Suppression of field population of *C. pomonella* by pilot release of sterile males

Pilot release of sterile males were conducted between July 8 and September 2, 2023, in Wumeifang Village, Jiuzhai Town, Gaizhou City, Yingkou City, Liaoning Province, China (40 ° 3 ′ 52 ”N, 122 ° 5 ′ 44 ”E). Prior to release, a central tree within the orchard was designated and marked, with sex traps positioned at 5-meter intervals in the cardinal directions of east, south, west, and north from the central tree. Sterile males were gently coated with fluorescent powder, distinguishing fumigated and unfumigated specimens by color before being released at 18:00. The control orchard, devoid of sterile male releases, was equipped with 5 trap sets, with trapped moth counts recorded at 10-day intervals post-release. Additionally, daily temperature and precipitation data were collected throughout the release period.

The five-point sampling method was employed in both the release orchard and control orchard to designate five points within the central and perimeter areas of the orchard. Around each of these points, five trees were meticulously selected and distinctly labeled. Prior to the release, a comprehensive examination of the total fruit count and the specific number of fruits was conducted. Subsequently, at intervals of 10 days, the quantity of infested fruits was meticulously documented, allowing for the calculation of the fruit infestation rate. This rate was determined by dividing the total number of infested fruits by the overall number of fruits examined, expressed as a percentage.

### 2.10 Data analysis

The statistical significance of various biological parameters of *C. pomonella*, encompassing metrics such as average emergence rate, longevity, fecundity, sterility rate, mating rate, as well as the flight characteristics of males including flight time, distance, and speed, along with gene expression in controlled laboratory settings and dispersal distance within the orchard, were meticulously analyzed utilizing the one-way analysis of variance (ANOVA) coupled with Duncan’s test (*P* ≤ 0.05) through SPSS 19 software (IBM Corporation, Armonk, NY, USA). Furthermore, significant difference in total number of moths trapped per trap, amount of insects trapped and fruit decay rate was analyzed through ANOVA with independent samples t-tests (\**P* < 0.05; \*\**P* < 0.01; \*\*\**P* < 0.001) using SPSS 19. The results were graphically represented using GraphPad Prism 5 (GraphPad Software, San Diego, CA, USA), with all values being expressed as mean ± standard error.

Additionally, the sterility rate of *C. pomonella* under varying X-ray dosage levels was modeled using a logistic curve in Origin 2019 software (Origin Lab Corporation, Northampton, MA 02115, USA).

## 3. RESULTS

### 3.1 Effect of different doses of X-ray irradiation on the biological parameters of *C. pomonella*

The effects of different doses of X-ray irradiation on the physiological traits of *C. pomonella* were explored in a series of experiments that examined the emergence rate, lifespan, reproductive capacity, and sterility of 8-day-old pupae exposed to varying levels of X-ray radiation. The results indicated that there was no statistically significant variance in the emergence rate of male moths *C. pomonella* within the range of 50-250 Gy compared to the control group. Notably, a reduction in the emergence rate of male *C. pomonella* moths was evident at 300 Gy (Figure 1a). However, the survival rate of moths arising from male pupae irradiated with various doses of radiation did not exhibit a significant difference compared to the control group (Figure 1b). Furthermore, a decrease in lifespan was evident with X-ray exposure surpassing 200 Gy when compared to the control group (Figure 1c). Fecundity levels remained steady across different radiation levels compared to the control group (Figure 1d). Additionally, an escalation in sterility rate was observed at doses below 250 Gy (Figure 1e), while a reduction in mating rate was noted at doses exceeding 200 Gy (Figure 1f).

**Figure 1.**
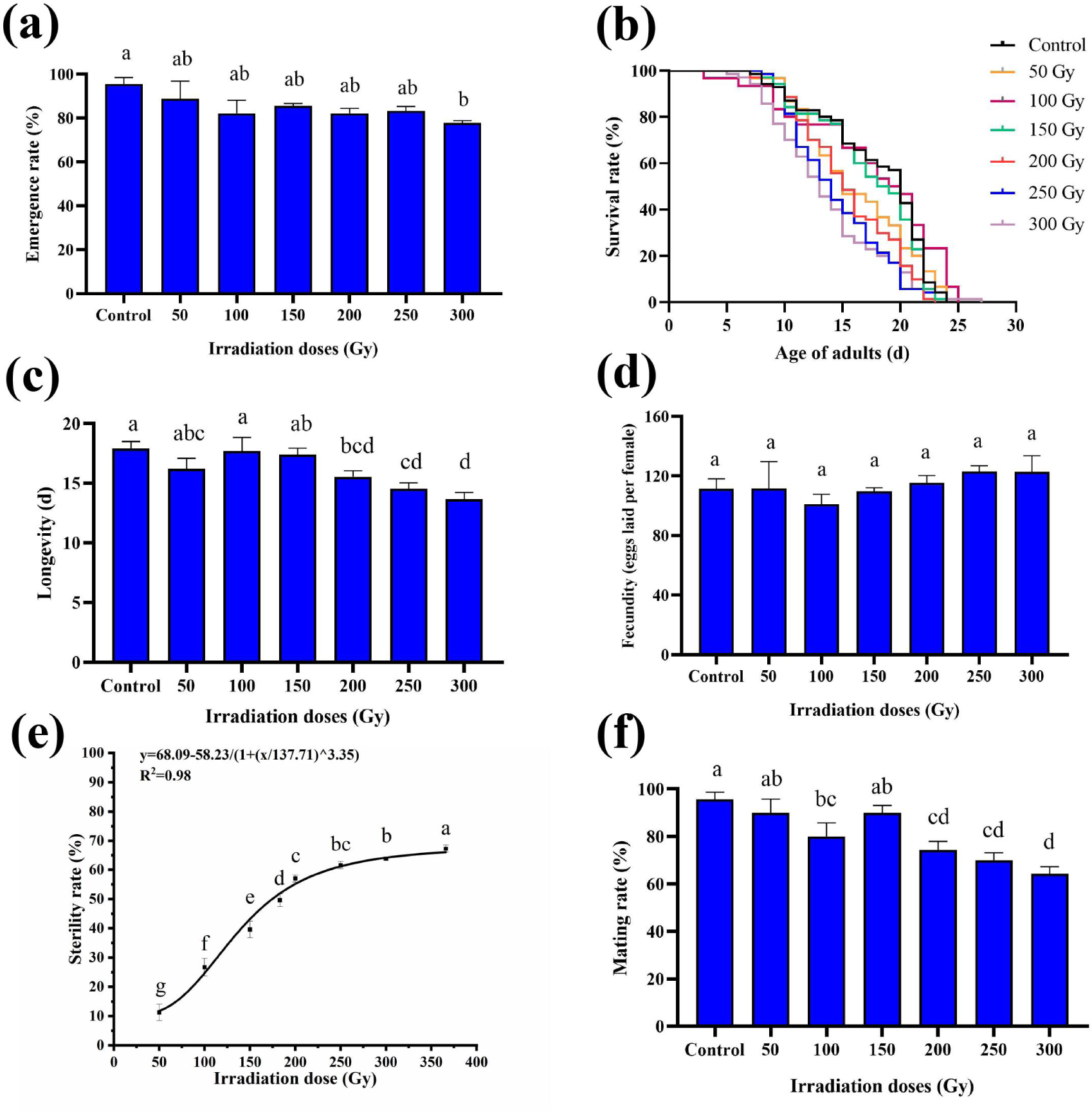
Effect of different doses of X-ray irradiation on the biological parameters of male *C. pomonella*. (a) Emergence rate; (b) Survival rate; (c) Longevity; (d) Fecundity (eggs laid per female); (e) Sterility rate; (f) Mating rate. Letters on the error bars indicate significant differences analyzed by the one-way analysis of variance (ANOVA) with Duncan’s test.

### 3. 2 Effect of 200 Gy X-ray irradiation on mating competitiveness of male *C. pomonella*

#### 3.2.1 Laboratory test

The hatching rate of eggs for SM:NM:NF at the 1:0:1 ratio was found to be 12.57 ± 0.51%, a value significantly lower compared to the ratios of 0:1:1 (72.96±7.31%) and 1:1:1 (64.18±3.95%). When male adults were combined with different ratios and matched with normal female adults, the expected hatching rate (E) was 42.77%. The competitive mating index (C) was determined to be 0.17, indicating that the mating competitiveness of male moths was reduced by 200 Gy of irradiation (Table 1).

**Table 1.** 200Gy X-ray irradiation on the mating competitiveness of *C. pomonella* adults.

| Experimental site | Matching ratio<br>(S♂: N♂: N♀) | Fecundity<br>(eggs laid per female) | Hatching rate/% |  | Competitive capacity |
| --- | --- | --- | --- | --- | --- |
|  |  |  | Observed value | Expected value |  |
| Indoor experiment | 0 : 10 : 10 | 116.20±19.97 a | 72.96±7.31 a |  |  |
|  | 10 : 0 : 10 | 103.75±4.42 a | 12.57±0.51 b |  |  |
|  | 10 : 10 : 10 | 106.07±5.80 a | 64.18±3.95 a | 42.77 | 0.17 |
| Field experiment | 0 : 10 : 10 | 57.45±4.09 a | 64.16±6.11 a |  |  |
|  | 10 : 0 : 10 | 60.13±10.60 a | 10.82±4.46 b |  |  |
|  | 10 : 10 : 10 | 54.26±6.81 a | 59.49±2.47 a | 37.49 | 0.096 |

Data are mean± SE. N: Number of unirradiated males; S: Number of irradiated males.

#### 3.2.2 Orchard trial

The hatching rate of eggs for the SM:NM:NF group at the 1:0:1 ratio was observed to be 10.82±4.46%, a significantly lower rate compared to the 0:1:1 ratio (64.16 ± 6.11%) and the 1:1:1 ratio (59.49 ± 2.47%). When male adults were introduced in various ratios and paired with normal female adults, the expected hatching value (E) was calculated to be 37.49%. Additionally, the competitive mating index (C) was determined to be 0.096, suggesting that a radiation dosage of 200 Gy diminished the mating competitiveness of male moths (Table 1).

### 3.3 Effects of different concentrations of linalool fumigation on biological parameters of *C. pomonella*

To elucidate the impact of varying concentrations of linalool fumigation treatments on the biological parameters of *C. pomonella*, experimental investigations were conducted to assess the survival duration, oviposition, egg hatching efficacy, and mating activity of male moths. The results revealed no notable disparities in the aforementioned parameters among male moths subjected to distinct levels of linalool fumigation in comparison to the control cohort, indicating that the application of linalool fumigation did not adversely affect the male moths (Figure 2).

**Figure 2.**
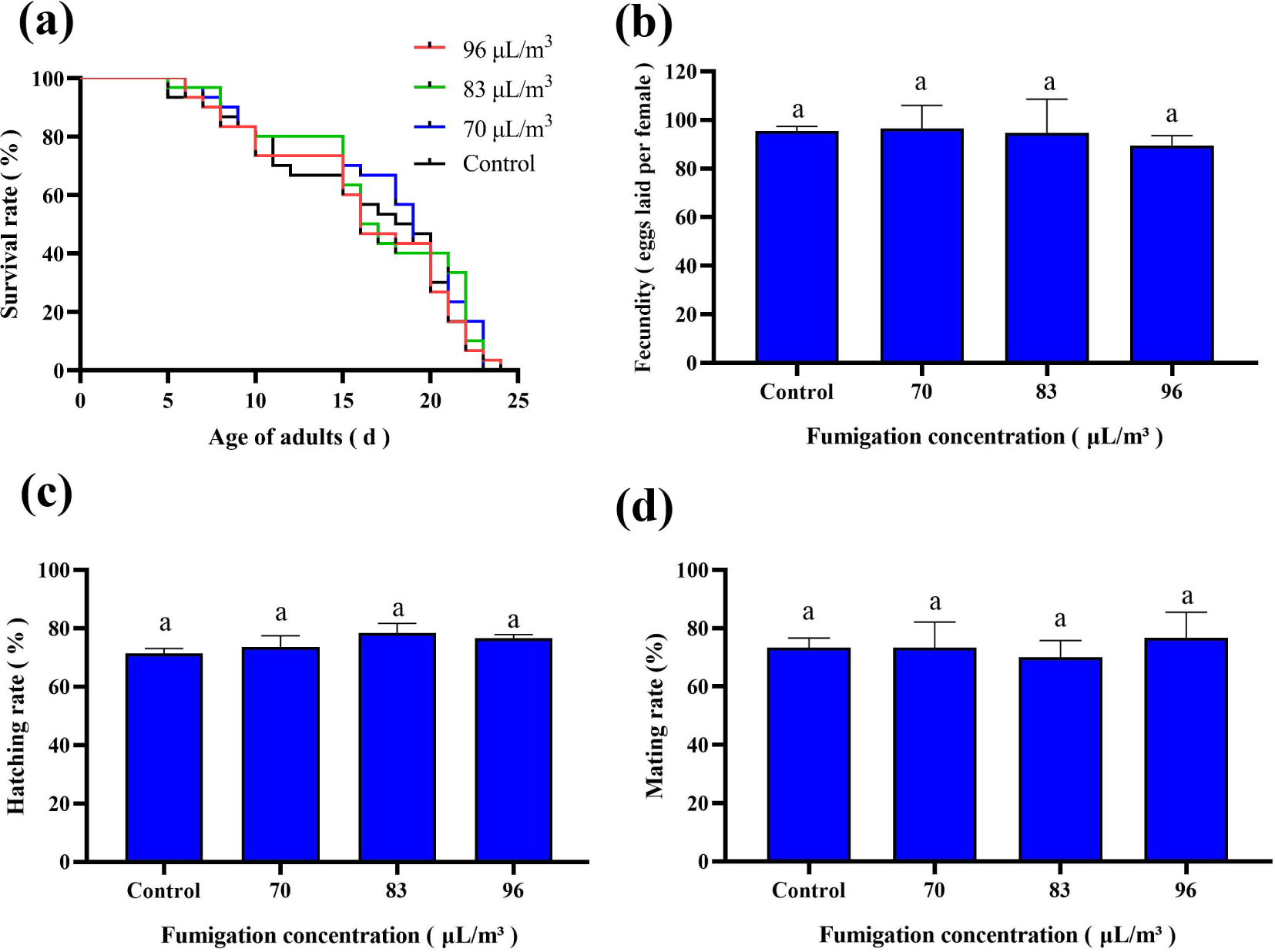
Effects of different concentrations of linalool fumigation on the biological parameters of male *C. pomonella*. (a) Survival rate; (b) Longevity; (c) Fecundity (eggs laid per female); (d) Hatching rate; (e) Mating rate.Letters on the error bars indicate significant differences analyzed by the one-way analysis of variance (ANOVA) with Duncan’s test.

### 3.4 Effects of different concentrations of linalool fumigation on mating competitiveness of *C. pomonella*

The hatching rate of eggs resulting from the pairing of fumigation-treated normal males with normal females were determined to be 73.63±3.86 percent, while the pairing of untreated normal males with normal females yielded a hatching rate of 71.42 ± 1.74%. When comparing the hatching rate of fumigation-treated normal males to the overall hatching rate, the value was found to be 72.55±0.33%. There was no significant difference observed in the hatchability of fecundity across three different mating combinations, with an average rate of 72.52%. Furthermore, the mating competitiveness index of *C. pomonella* normal males that were exposed to linalool fumigation at a concentration of 70 μ L/m ³ was calculated to be 1.06, suggesting an enhancement in the mating competitiveness of the males (Table 2).

**Table 2.**
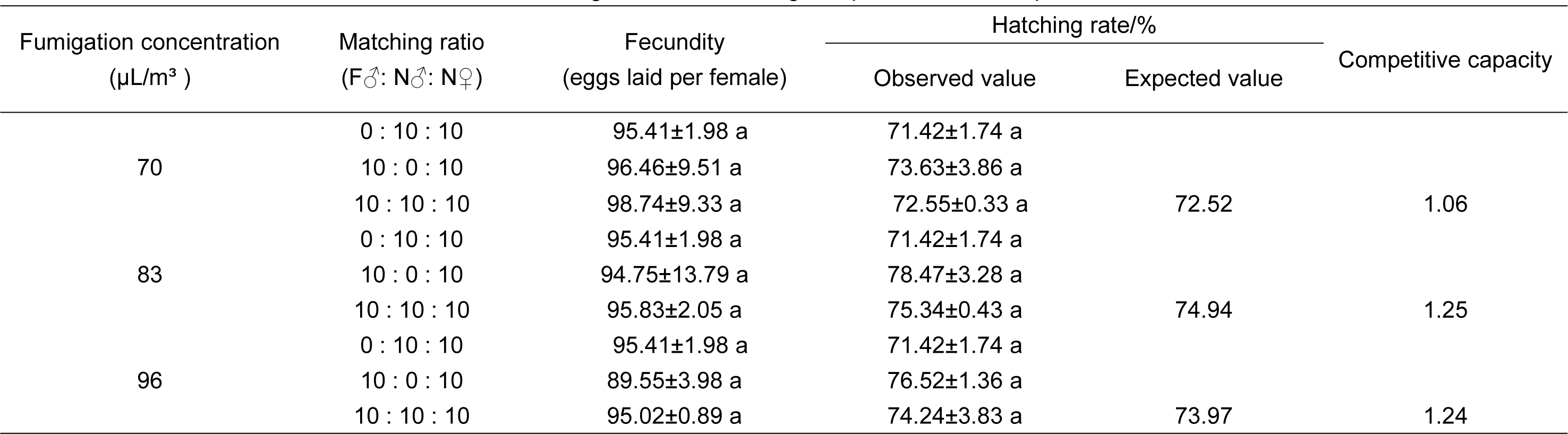
Effect of different concentrations of linalool fumigation on the mating competitiveness of *C. pomonella* males.

The percentage of eggs hatched when 83 μL/m³ of linalool was used to fumigate normal males paired with normal females was recorded at 78.47 ± 3.28%. For normal males paired with normal females, the hatching rate was slightly lower at 71.42 ± 1.74%. When fumigated normal males were paired with normal females, the hatching rate was 75.34 ± 0.43%. There was no significant difference in the hatching rate of eggs produced by the three different mating combinations, all averaging around the expected value of 74.94. The mating competitiveness index of males treated with linalool at 83 μ L/m ³ was 1.25, indicating an improvement in mating competitiveness compared to those treated at 70 μL/m³ (Table 2).

The egg hatching rate of 96 μL/m³ linalool-exposed normal male moths when paired with normal females was determined to be 76.52±1.36%, while that of normal males paired with normal females was 71.42±1.74%, and the hatching rate of fumigated normal males when paired with other males and females was 74.24 ± 3.83%. These results indicate that there were no significant differences in hatching rates among the three mating combinations studied, all of which were close to the anticipated value of 73.97%. Furthermore, the mating competitiveness index of males exposed to linalool at 96 μ L/m ³ was calculated to be 1.24, suggesting a reduced mating competitiveness compared to males exposed to linalool at 83 μL/m³ (Table 2).

### 3.5 Effect of linalool fumigation on mating competitiveness of sterilized male *C. pomonella* moths

#### 3.5.1 Laboratory test

The results revealed that male moths exposed to 200 Gy X-rays exhibited notably reduced hatching and mating frequencies in comparison to the untreated group. Notably, there were no substantial variations in any of the outcomes between the fumigated and non-fumigated sterilized males, indicating that the linalool fumigation did not impact the reproductive characteristics of sterilized males (Figure 3).

**Figure 3.**
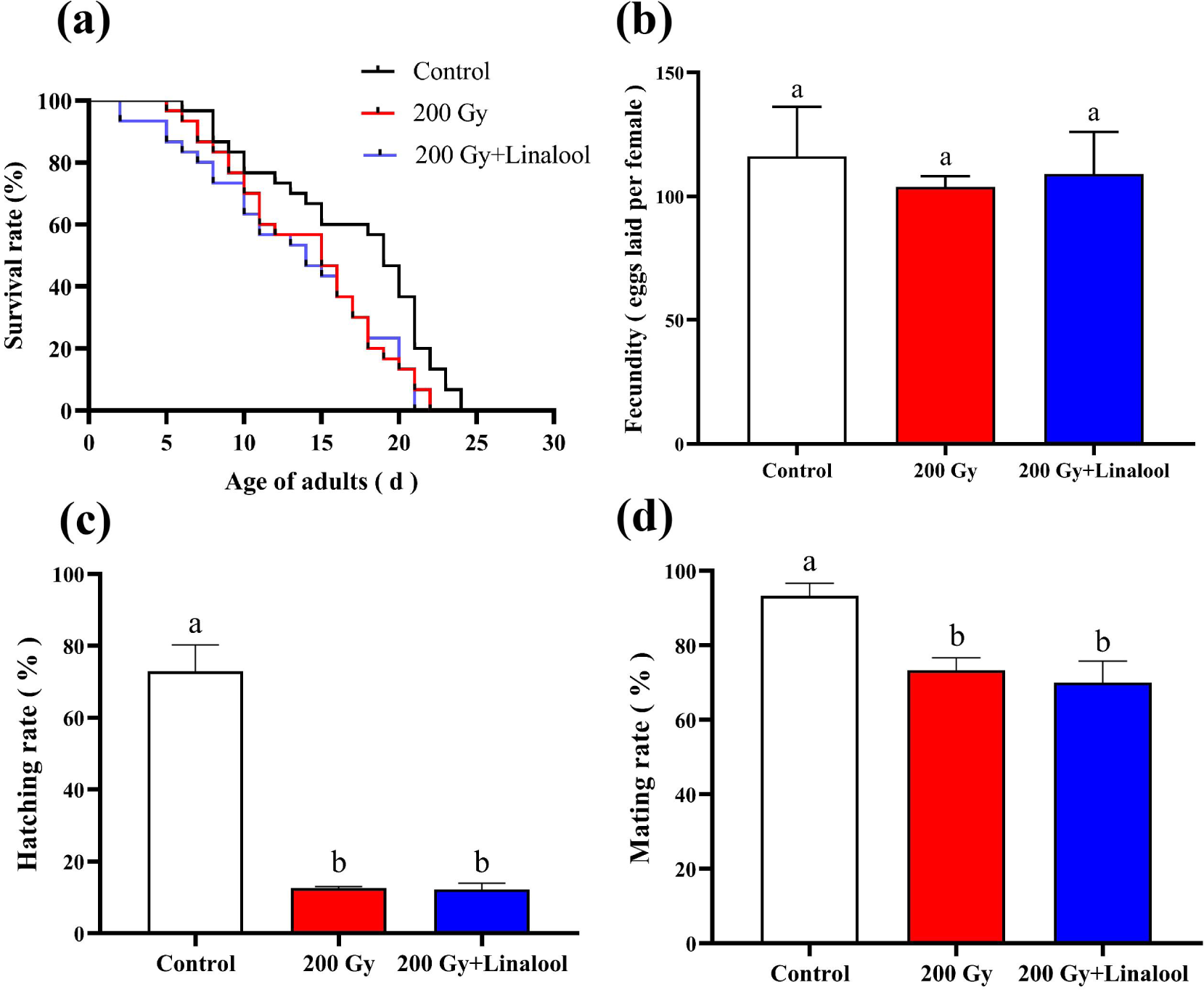
Effects of different concentrations of linalool fumigation on the biological parameters of 200 Gy X-ray sterilized male *C. pomonella* (laboratory experiment). (a) Survival rate; (b) Longevity; (c) Fecundity (eggs laid per female); (d) Hatching rate; (e) Mating rate. Letters on the error bars indicate significant differences analyzed by the one-way analysis of variance (ANOVA) with Duncan’s test.

**Figure 4.**
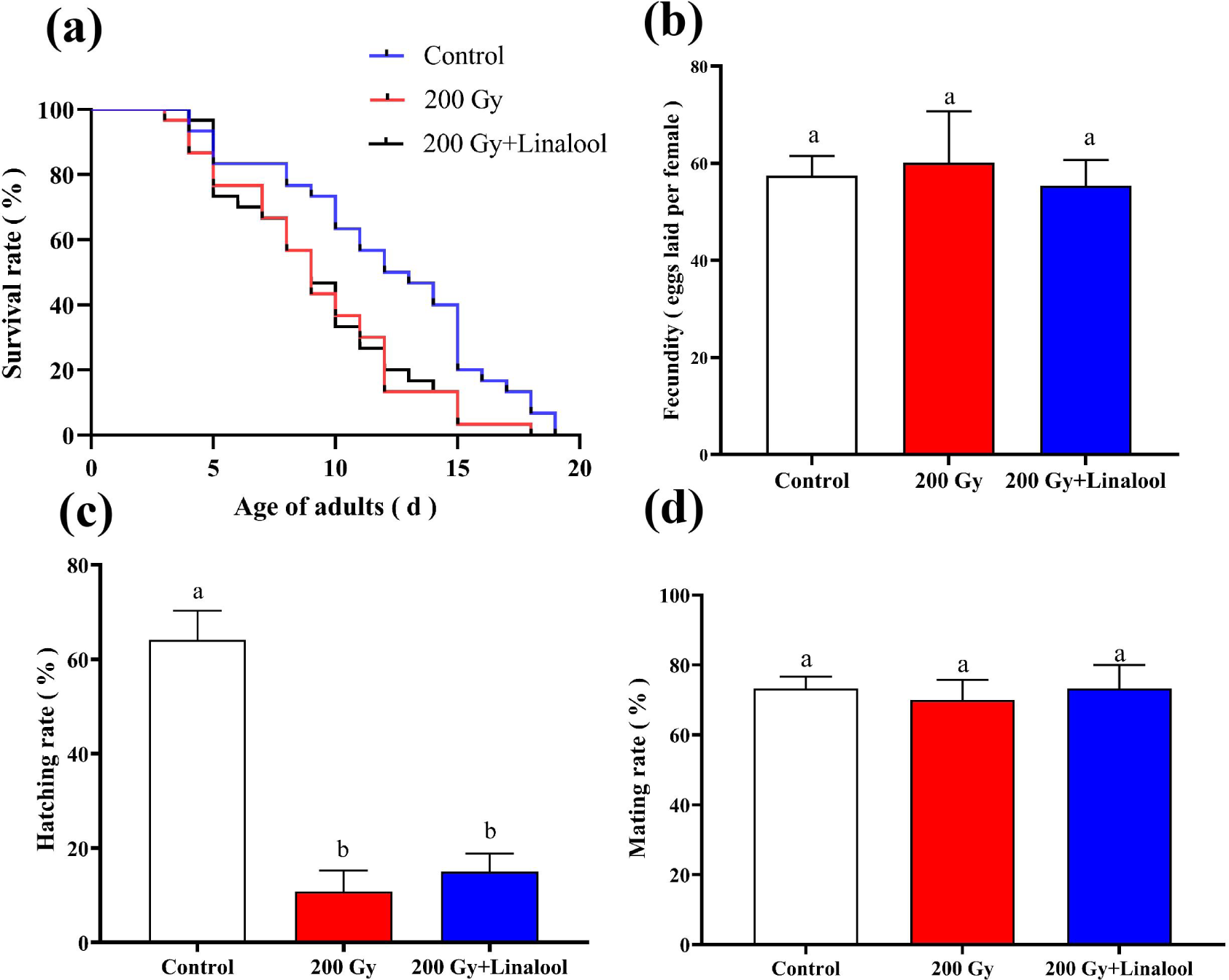
Effect of different concentrations of linalool fumigation on the biological parameters of 200 Gy X-ray sterilized male *C. pomonella* (field experiment). (a) Survival rate; (b) Longevity; (c) Fecundity (eggs laid per female); (d) Hatching rate; (e) Mating rate. Letters on the error bars indicate significant differences analyzed by the one-way analysis of variance (ANOVA) with Duncan’s test.

Specifically, the hatching rate of linalool-fumigated irradiated sterilized males that were paired with normal females was 13.44± 2.33%, while that of irradiated sterilized males paired with normal females stood at 10.97±2.41%.

Moreover, the hatching rate of fumigation-treated irradiated sterilized males when paired with females was 12.27 ± 1.98%, suggesting no significant variance in the hatching rate of eggs produced by the three different pairings (F = 0.305, df = 8, P = 0.748), with an expected value of 12.21%. The mating competition index of 1.11 for the irradiated sterilized males of moths fumigated with linalool indicated an enhancement in the mating competitiveness of the irradiated sterilized males due to linalool fumigation (Table 3).

**Table 3.** Effect of different concentrations of linalool fumigation on the mating competitiveness of 200 Gy X-ray sterilized *C. pomonella* males.

| Experimental site | Matching ratio<br>(F♂: N♂: N♀) | Fecundity(eggs laid per<br>female) | Hatching rate/% |  | Competitive capacity |
| --- | --- | --- | --- | --- | --- |
|  |  |  | Observed value | Expected value |  |
| Indoor experiment | 0 : 10 : 10 | 106.54 ±5.71 a | 10.97±2.41 a |  |  |
|  | 10 : 0 : 10 | 103.30± 12.00 a | 13.44±2.33 a |  |  |
|  | 10 : 10 : 10 | 94.35± 11.43 a | 12.27±1.98 a | 12.21 | 1.11 |
| Field experiment | 0 : 10 : 10 | 60.13±10.60 a | 10.82±4.46 a |  |  |
|  | 10 : 0 : 10 | 55.40±5.28 a | 14.98±3.89a |  |  |
|  | 10 : 10 : 10 | 61.66±4.09 a | 12.84±1.98 a | 12.90 | 1.06 |

#### 3.5.2 Orchard trial

The results of the field experiment revealed a significant decrease in the longevity and fecundity of sterile males of *C. pomonella* compared to the control group. Interestingly, there was no notable variance between the biological parameters of linalool-fumigated sterile males and non-fumigated ones, suggesting that the fumigation treatments did not impact these parameters within the field environment (Figure 3).

When linalool-fumigated irradiated sterile males were paired with normal females, the hatchability was observed to be 14.98±3.89%, while it was 10.82 ±4.46% for *C. pomonella* irradiated sterile males mated with normal females.

Moreover, the hatchability of fumigation-treated irradiated sterile males when paired with females stood at 12.84±1.98%. These findings indicate that there was no significant difference in hatchability and fecundity among the three different combinations (F = 0.249, df = 8, P = 0.756), with an average value of 12.90%. The mating competition index of 1.06 for the linalool-fumigated irradiated sterile males of *C. pomonella* suggests that linalool fumigation enhanced the mating competitiveness of the irradiated sterile males (Table 3).

### 3.6 Effects of linalool fumigation on the flight ability of sterile *C. pomonella* moths

The flight duration of 3-day-old male adults, sterile males, and fumigation treated males was 53.82±7.17, 5.61±2.36, and 10.18±3.46 min, respectively.

While no significant disparity was observed in the flight duration between sterile males and fumigation treated males, both groups exhibited significantly shorter flight times compared to the control group (F = 30.820, df = 44, P < 0.001). Moreover, there were no significant differences in flight distance or flight speed among the treatment groups when compared to the control group, (Figure 5).

**Figure 5.**
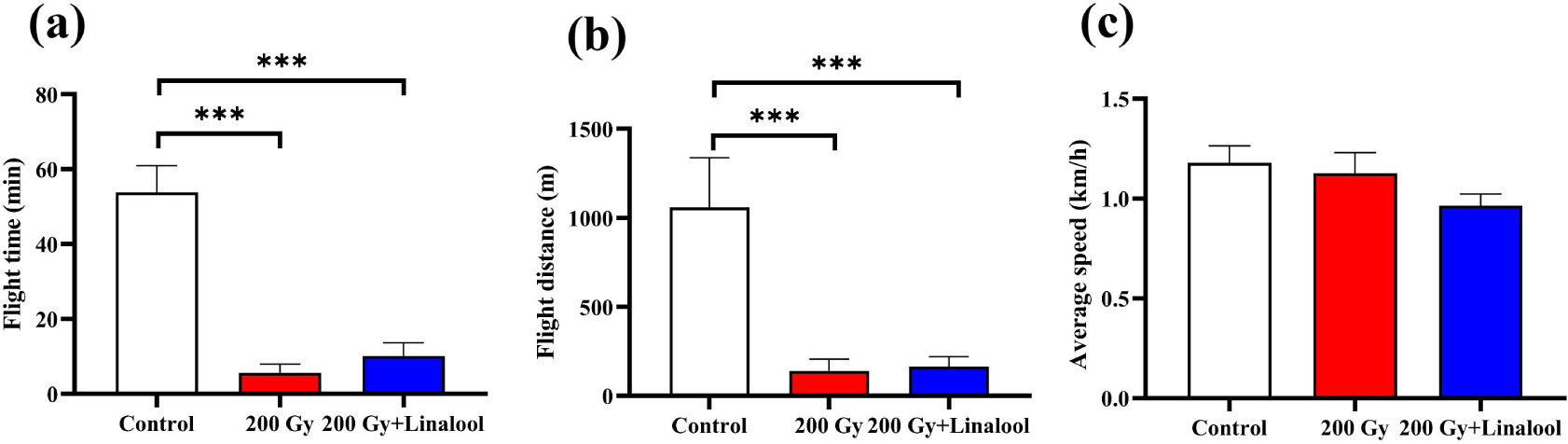
Effects of X-ray irradiation on the flight ability of male adults of *C. pomonella.* (a) Flight duration; (b) Flight distance; (c) Average flight speed. Asterisk (*) on the error bars indicate significant differences analyzed by the one-way analysis of variance (ANOVA) with independent samples t-tests (*, *P* < 0.05; **, *P* < 0.01; ***, *P* < 0.001).

### 3.7 Effects of linalool fumigation on the expression levels of sex pheromone receptor-related genes in sterile *C. pomonella* moths

RT-qPCR results showed that *CpomOR1, CpomOR2a, CpomOR3a, CpomOR3b, CpomOR5, CpomOR6a, CpomOR7, CpomGOBP1,* and *CpomGOBP2* genes were significantly reduced in *C. pomonella* male moths that emerged from pupae irradiated with 200 Gy X-ray (Figure 6).

**Figure 6.**
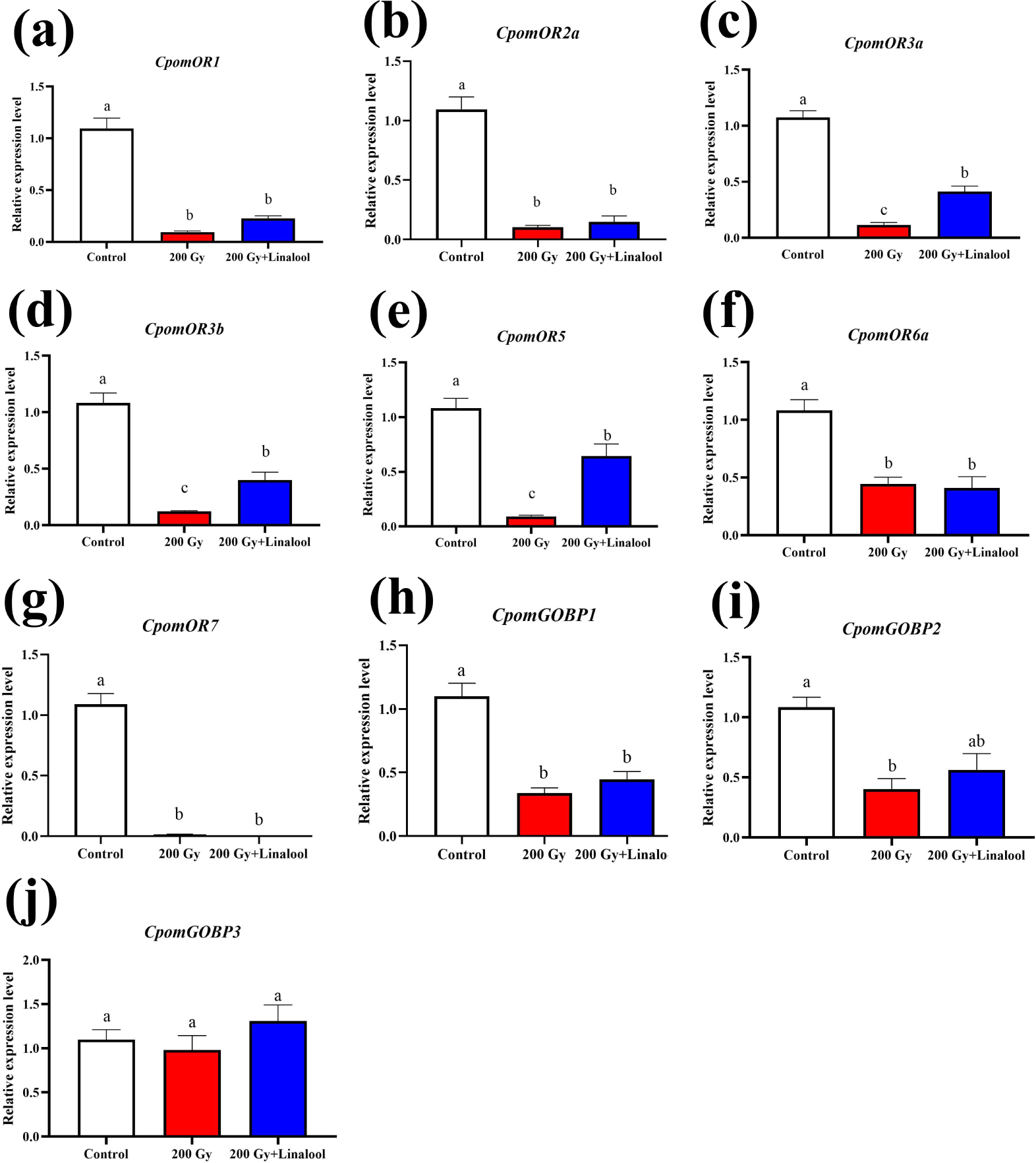
Effect of X-ray irradiation on the relative expression levels of the pheromone recognition-related genes in male adults of *C. pomonell*a. (a) *CpomOR1*; (b) *CpomOR2*a; (c) *CpomOR3*a; (d) *CpomOR3b*; (e) *CpomOR5*; (f) *CpomOR6a*; (g) *CpomOR*7; (h) *CpomGOBP*1; (i) *CpomGOBP2*; (j) *CpomGOBP3*. The 8-day-old male pupae (1 day before emergence) were exposed to 200 Gy of X-ray, and the expression levels of pheromone receptor-related genes in 3-day-old male adults were analyzed. Letters on the error bars indicate significant differences analyzed by the one-way analysis of variance (ANOVA) with Duncan’s test.

The expression levels of *CpomOR3a, CpomOR3b,* and *CpomOR5* were significantly rebounded after linalool fumigation (Figure 6).

### 3.8 Suppression of field population of *C. pomonella* by pilot release of sterile males

A total of 5342 sterile males were released, with 2735 of them being released from linalool fumigated moths. Among these, 60.44±4.48% of males flew out normally, while 6.83 ± 1.51% fell to the ground post take-off, leaving 39.56±4.48% unable to take off normally (Table S2). The percentage of males falling to the ground was 59.84±6.86%. Additionally, 2607 sterile males of *C. pomonella* without linalool fumigation were also released, where 62.17±4.25% flew out normally, 17.07 ± 2.86% fell to the ground after take-off, and 37.83 ± 4.25% couldn’t take off normally. Notably, 78.19±6.36% of males picked with tweezers failed to take off and fell to the ground (Table S3).

Throughout the experiment, the ambient temperature at the experimental site exhibited minimal variations, with a singular precipitation event exceeding 100 mm (Figure S9). Results revealed that, at a 5-meter distance, the recapture of sterile male moths in traps averaged 12.25 ± 3.07, whereas the recapture of fumigation-treated sterile males was 8.75 ± 3.16. Similarly, at distances of 10 m, the recaptures were 10.00 ± 1.78 and 9.75 ± 1.18, respectively. At 15 m, the retrievals were 8.50 ± 1.55 and 6.50 ± 1.19, respectively. Finally, at a distance of 20 m, the retrievals were 0±0 and 1.50± 0.65, respectively. The results indicated that there were no statistically significant variations in the recaptures of sterile males and fumigation-treated sterile males of *C. pomonella* at the 10 and 20 m distances at a significance level of 0.05 (P > 0.05). However, notable differences were noted in recaptures at 5 and 15 meters, with statistically significant variances observed at the 0.05 level (P < 0.05) (Figure S10).

The deployment of *C. pomonella* sterile males over a period of two months resulted in a notable decline in fruit infestation within the treated orchards, registering at 24.87 ± 2.17%. This rate was significantly lower (*P* < 0.05) in contrast to the control orchard where infestation levels reached 34.01 ± 2.87%. These findings provide evidence that the introduction of sterile *C. pomonella* males effectively mitigated fruit infestation occurrences, thereby diminishing both the incidence of infested fruits and the overall infestation rate within the orchard (Figure 7C).

**Figure 7.**
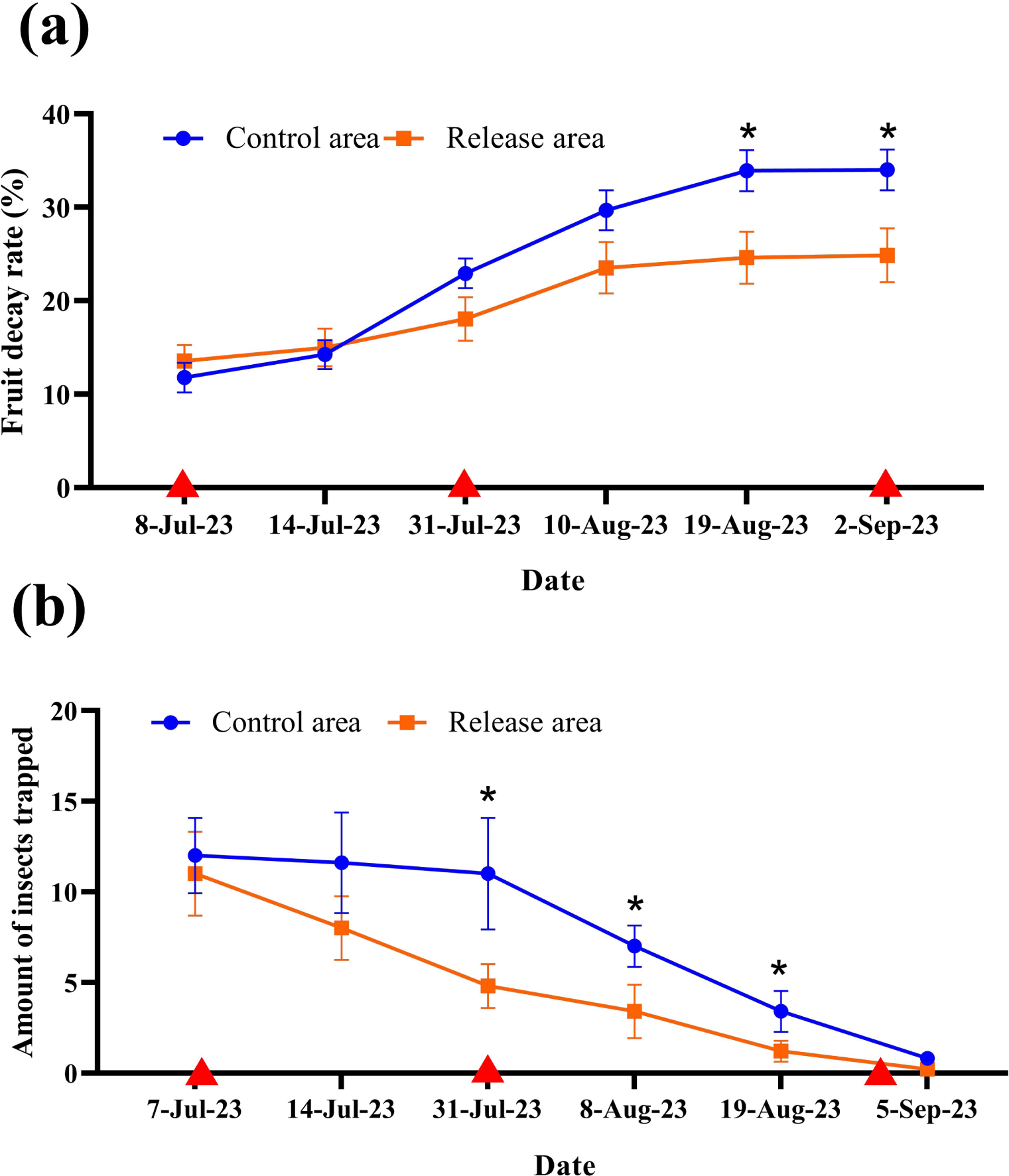
Suppression of field population of *C. pomonella* by pilot release of sterile male moths in an apple orchard. (a) Survey of fruit decay rate in control and release areas. The red triangles represent the date of release. (b) Number of sterile males recaptured by traps at different distances. (*) on the error bars indicate significant differences analyzed by the one-way analysis of variance (ANOVA) with independent samples t-tests (*, *P* < 0.05; **, *P* < 0.01; ***, *P* < 0.001).

The captures of wild male moths did not exhibit a notable variance between the release area and the control from July 7 to July 14, 2023. Nonetheless, subsequent to July 31, 2023, a noteworthy decline in captures within the release area was observed, presenting a significant contrast with the control group (P < 0.05). Based on these findings, it is evident that the introduction of sterile male *C. pomonella* moths in the natural environment substantially reduces the population of wild male moths (Figure 7d).

## 4 DISCUSSION

The radiation dose is a critical factor in achieving the desired sterility level of insects in the SIT program.^8, 27^ In this study, we demonstrated that the biological traits of male *C. pomonella* varied in their decline with increasing levels of irradiation. This observation is consistent with a previous study on *Glossina palpalis* tsetse flies, which showed that higher radiation doses reduced the fecundity of females.^28^ Additionally, we observed a significant decrease in the mating rate of sterile male *C. pomonella* with normal females when exposed to radiation above 200 Gy, mirroring the findings in *G. palpalis*.^28^ The OK SIR program, designed to control *C. pomonella* populations, initially utilized 250 Gy of ^60^Co-γ irradiation, resulting in male sterility rates of 85% to 92%.^29^ Since 2014, a consistent dose of 200 Gy has been employed to balance sterility levels and mating competitiveness.^30^ To optimize the trade- off between male sterility and minimize the negative effects of radiation on the insects, it is recommended to use 200 Gy as the optimal dose for inducing sterility in males *C. pomonella*.

The mating competitiveness of sterile insects is crucial for their ability to effectively compete with wild males for mating opportunities with females.^31,32^ Previous research has shown that inadequate dosages may result in incomplete sterilization, while excessive irradiation can reduce the mating competitiveness of released insects compared to wild counterparts, thus affecting the success of SIT programs^.33^ Therefore, maintaining a delicate balance between mating competitiveness and sterility is essential during copulatory interactions with wild female insects.^34^ Evaluating the optimal irradiation dose is critical to assessing the mating competitiveness of male insects bred for this purpose.^33^ However, reduced mating competitiveness is commonly observed in irradiated *C. pomonella* populations.^7^ Jiang et al.^35, 36^ also reported a significant decrease in the mating competitiveness of male *Spodoptera frugiperda* following exposure to 250 Gy of X-ray irradiation. In our recent studies, male *C. pomonella* pupae irradiated with 366 Gy of X-rays exhibited mating competitiveness indices of 0.001 in laboratory^13^ and 0.0088 in orchards.^37^ In this study, when subjected to 200 Gy of X-ray irradiation, the mating competitiveness index of sterilized male *C. pomonella* decreased, indicating a decline in mating competitiveness post-irradiation. Nevertheless, compared to the impact of 366 Gy irradiation, the mating competitiveness index of *C. pomonella* males following 200 Gy irradiation increased to 0.17 in laboratory conditions and 0.096 in orchards, suggesting that an optimized irradiation dose can enhance the mating competitiveness of sterile insects to a certain degree. However, the mating competitiveness of *C. pomonella* males post-exposure to 200 Gy irradiation persists at a sub-optimal level. Therefore, it is imperative to prioritize efforts aimed at improving the mating competitiveness of sterile males in future studies.

The diminished mating competitiveness evident in irradiated insects may stem from the compromised olfactory system in male insects, leading to a reduced ability to perceive female sex pheromones ^38^. To investigate this hypothesis, linalool, a volatile compound found in host plants known to enhance the attraction of *C. capitata* moths^16^, was utilized to fumigate male *C. pomonella* moths. Although linalool fumigation did not affect longevity, fecundity, hatching rates, or overall mating performance, it did enhance the mating competitiveness of male *C. pomonella* moths. Another investigation on *C. capitata* demonstrated that the mating competitiveness of normal male adults and mass-reared irradiated males increased subsequent to fumigation with ginger root oil (GRO), compared to non-fumigated adults. Notably, GRO fumigation did not have any notable impacts on the body weight or longevity of irradiated mass-reared males.^39^ These results suggest that linalool fumigation might potentially mitigate male sterility induced by X-ray irradiation, possibly due to enhanced male sex pheromone communication in the antennae of male *C. pomonella* moths, as observed in *C. capitata* post ginger root oil fumigation.^40^ Furthermore, we revealed a decrease in gene expression in *C. pomonella* males’ antenna following 200 Gy of X-ray irradiation, suggesting a reduced sex pheromones response in male moths. Conversely, an increase in the expression of *CpomOR3a*, *CpomOR3b*, and *CpomOR5* genes was observed after linalool fumigation, enhancing the recognition of sex pheromones. These findings support the efficacy of linalool fumigation in mitigating male sterility induced by X-ray irradiation, likely through the upregulation of essential chemosensory genes and the activation of pheromone recognition mechanisms in *C. pomonella*. Further confirmation through behavioral experiments, comparative genomic and transcriptomic analyses, and molecular biology experiments is required to determine if the reduced mating competitiveness of irradiated insects is due to dysfunction of the olfactory system and the involvement of additional chemosensory genes in the recognition of pheromones in male adults.

Previous studies have demonstrated that the release of *C. pomonella* males into the pear orchard following exposure to 366 Gy X-ray did not affect the wild populations or the fruit decay rate.^13^ In this study, we carried out a small-scale trial in different fields utilizing the SIT to manage *C. pomonella* populations. Our study demonstrated that by releasing three successive batches of sterile male *C. pomonella* moths irradiated at 200 Gy, along with linalool fumigation, in the specified orchard area, there was a substantial decrease in *C. pomonella* infestations in the apple crop. This reduction was evident through significantly lower infestation rates compared to the control orchard. Additionally, field observations indicated a notable decrease in the recapture rate of wild *C. pomonella* moths, aligning closely with the findings of Huang *et al*.^4^, in comparison to the control group. These results suggest that the utilization of sterile males effectively suppressed the wild *C. pomonella* population in the orchards, consequently reducing fruit damage. The findings highlight an enhancement in the field application of the SIT for *C. pomonella* control after optimizing the irradiation dose.

In summary, we have identified the optimal sterilizing dose of X-ray irradiation for *C. pomonella* and introduced a plant volatiles fumigation approach to enhance the mating competitiveness of sterilized males. Additionally, we investigated the effects of irradiation and fumigation on the expression of genes related to pheromone recognition in the male antennal. Notably, our study demonstrated the successful reduction of the *C. pomonella* population in the orchards through repeated releases of sterilized males in conjunction with linalool fumigation on a pilot scale. These findings offer valuable insights for the implementation of the SIT as a strategy of controlling *C. pomonella*.

## Acknowledgments

This research was funded by the National Key R&D Program of China (2021YFD1400200, 2023YFD1400704) and the IAEA TC program (CPR5027).

## Author Contributions Statement

Sheng-Wang Huang: software, methodology, investigation, formal analysis, writing original draft, Peng-Cheng Wang: software, methodology, investigation, formal analysis Yan Wang: software, review and editing. Jie-Qiong: formal analysis, conceptualization. Ping Gao: writing, review, and editing, resources. Xue-Qing Yang: supervision, resources, project administration, funding acquisition, conceptualization, writing, review, and editing.

## Supporting information

**Table S1.**
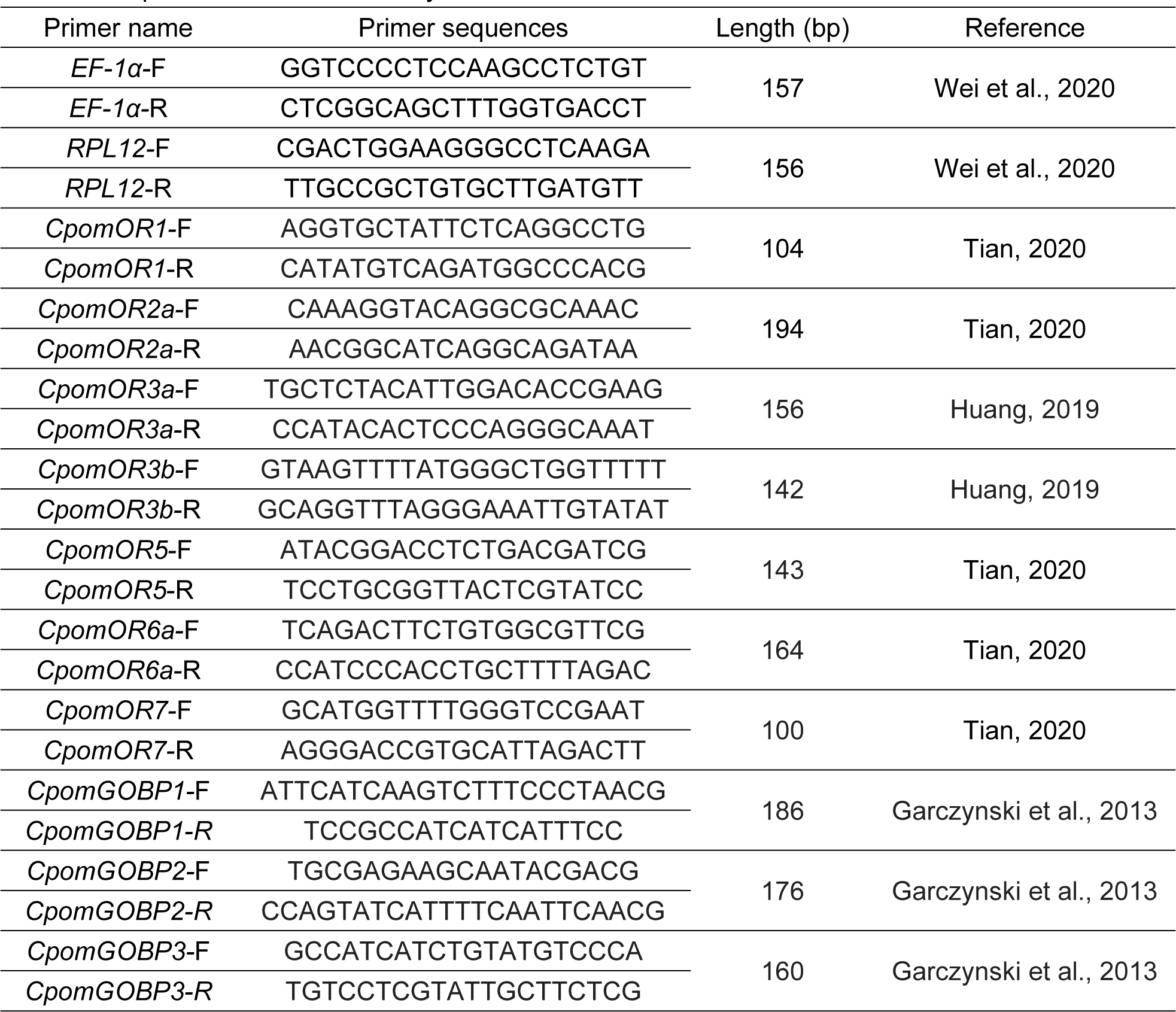
The primers used in this study.

**Table S2.** Flight status of released sterile males in the field.

|  | Total released | Normal flight out<br>(%) | Abnormal flight out<br>(%) |
| --- | --- | --- | --- |
| Sterile males exposed to<br>linalool | 2735 | 60.44±4.48 | 39.56±4.48 |
| Sterile males | 2607 | 62.17±4.25 | 37.83±4.25* |
| Total released | 5342 |  |  |

**Table S3.** Percentage of sterile males that can normal flight out but landed and abnormal flight out with landed after pick offs in the field.

|  | Normal flight out but<br>landed(%) | Abnormal flight out with<br>landed after pick offs (%) |
| --- | --- | --- |
| Sterile males exposed to<br>linalool | 6.83±1.51 | 59.84±6.86 |
| Sterile males | 17.07±2.86* | 78.19±6.36 |

**Figure S1.**
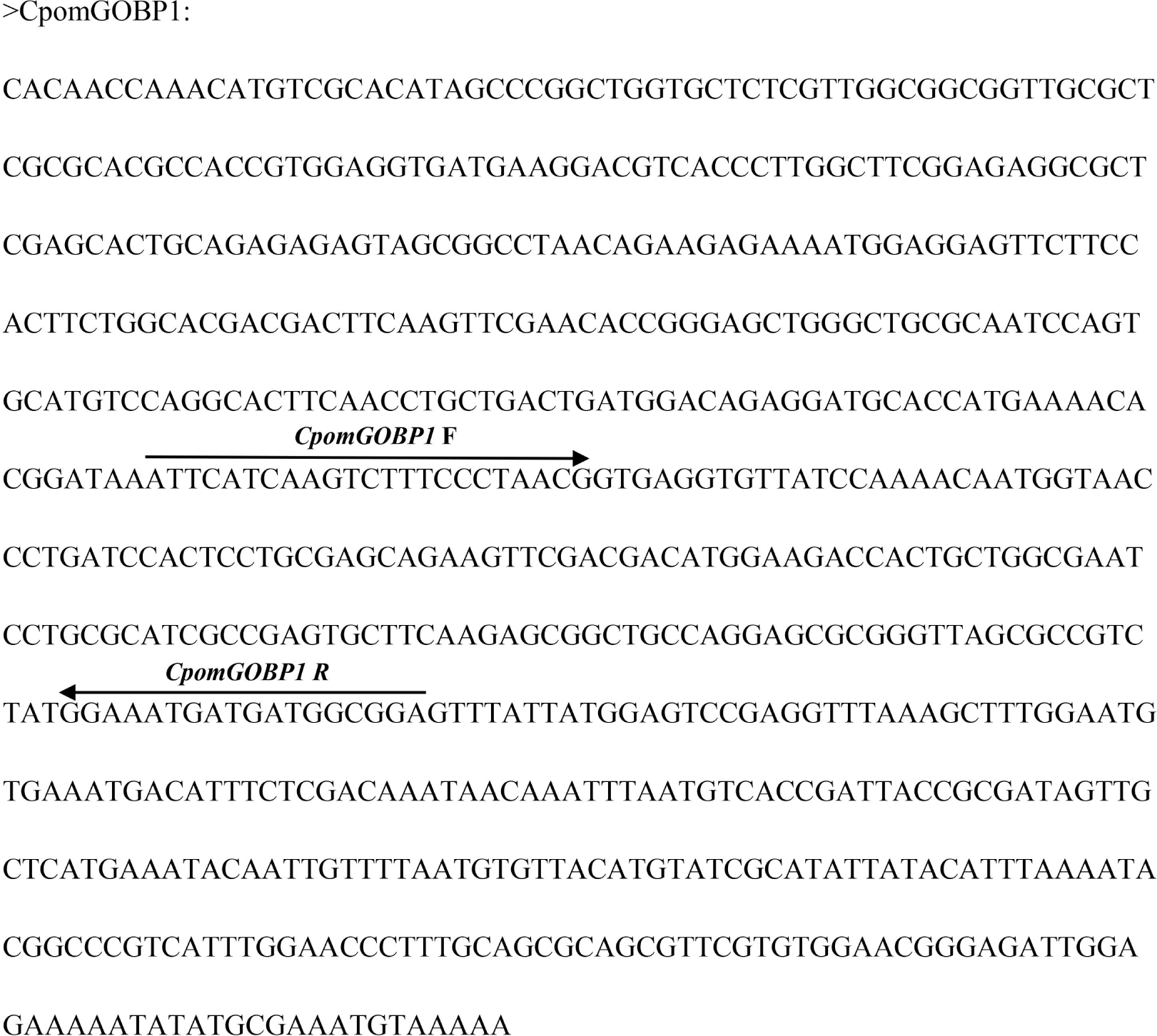
The cDNA sequence of CpomGOBP1.The location of the primer sequence is indicated by an arrow.

**Figure S2.**
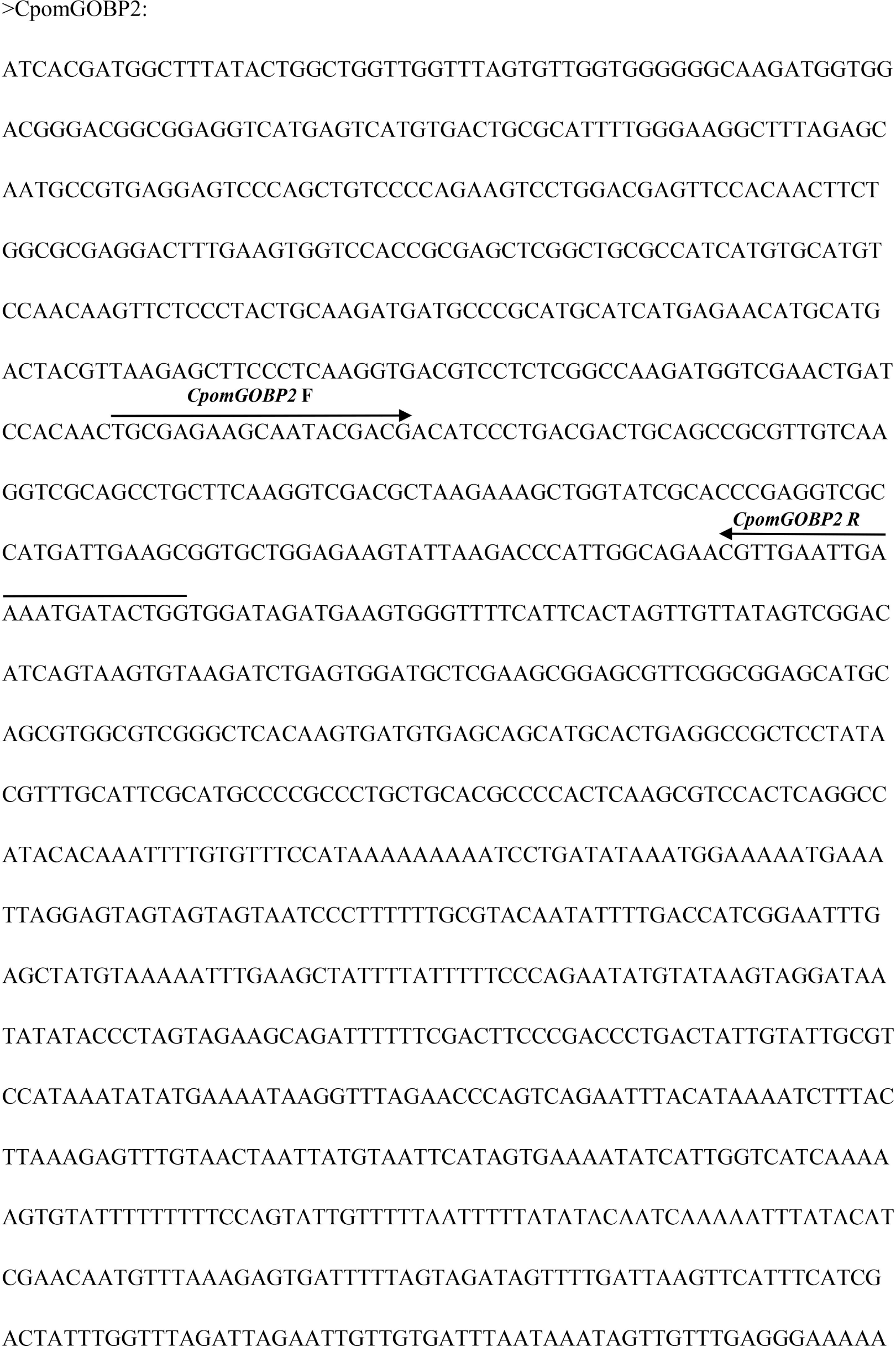
The cDNA sequence of CpomGOBP1.The location of the primer sequence is indicated by an arrow.

**Figure S3.**
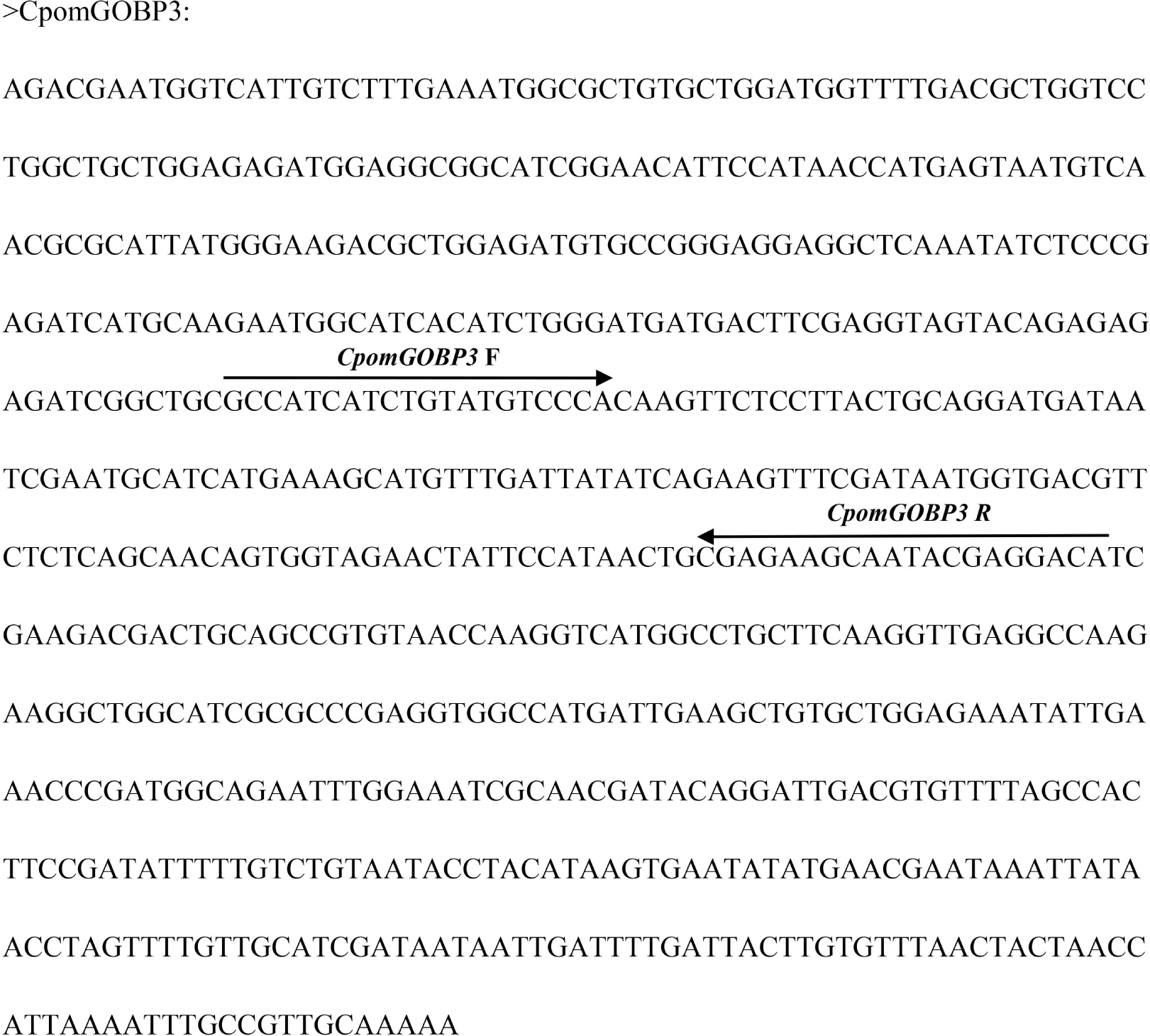
The cDNA sequence of CpomGOBP3.The location of the primer sequence is indicated by an arrow.

**Figure S4.**
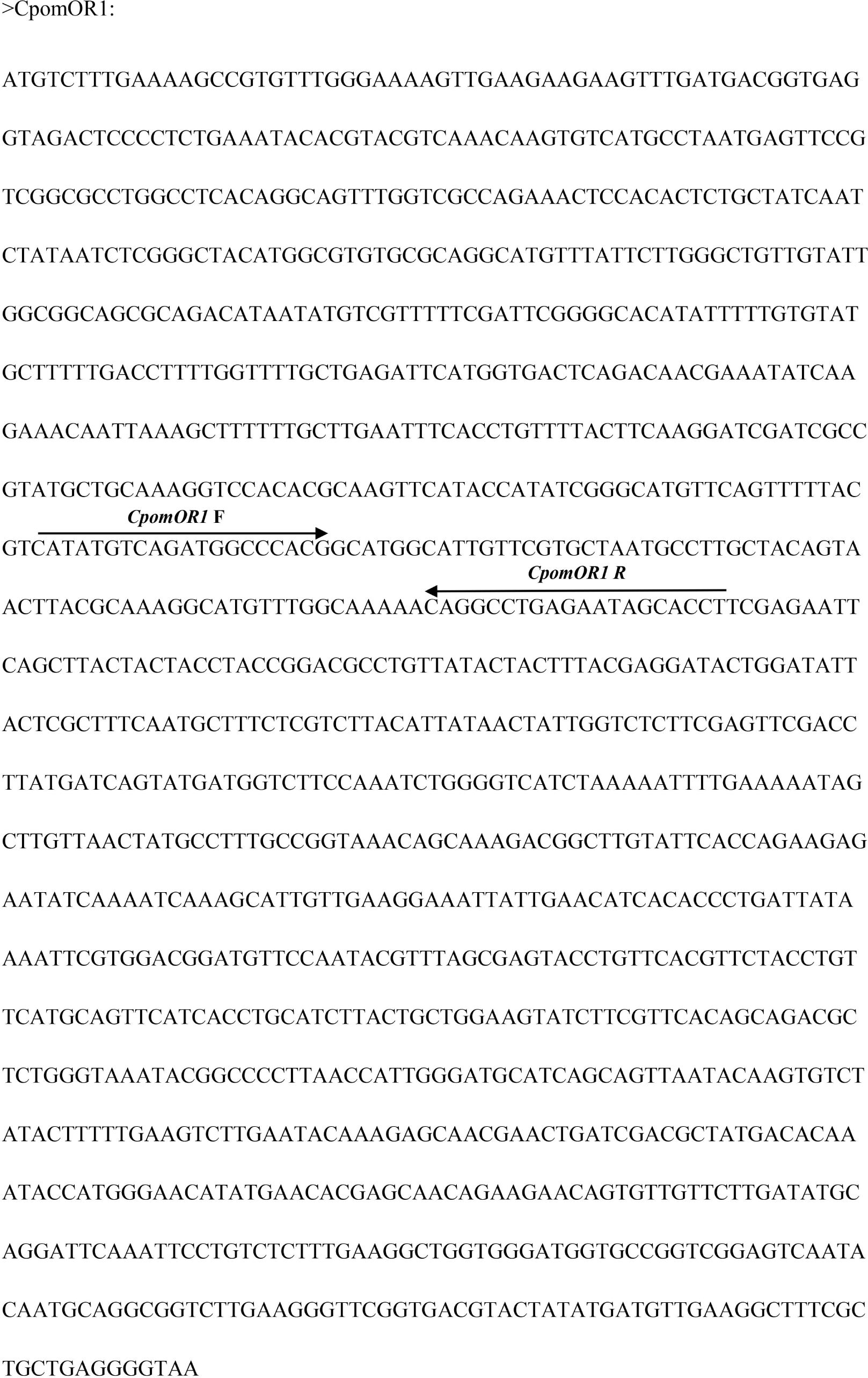
The cDNA sequence of CpomOR1.The location of the primer sequence is indicated by an arrow.

**Figure S5.**
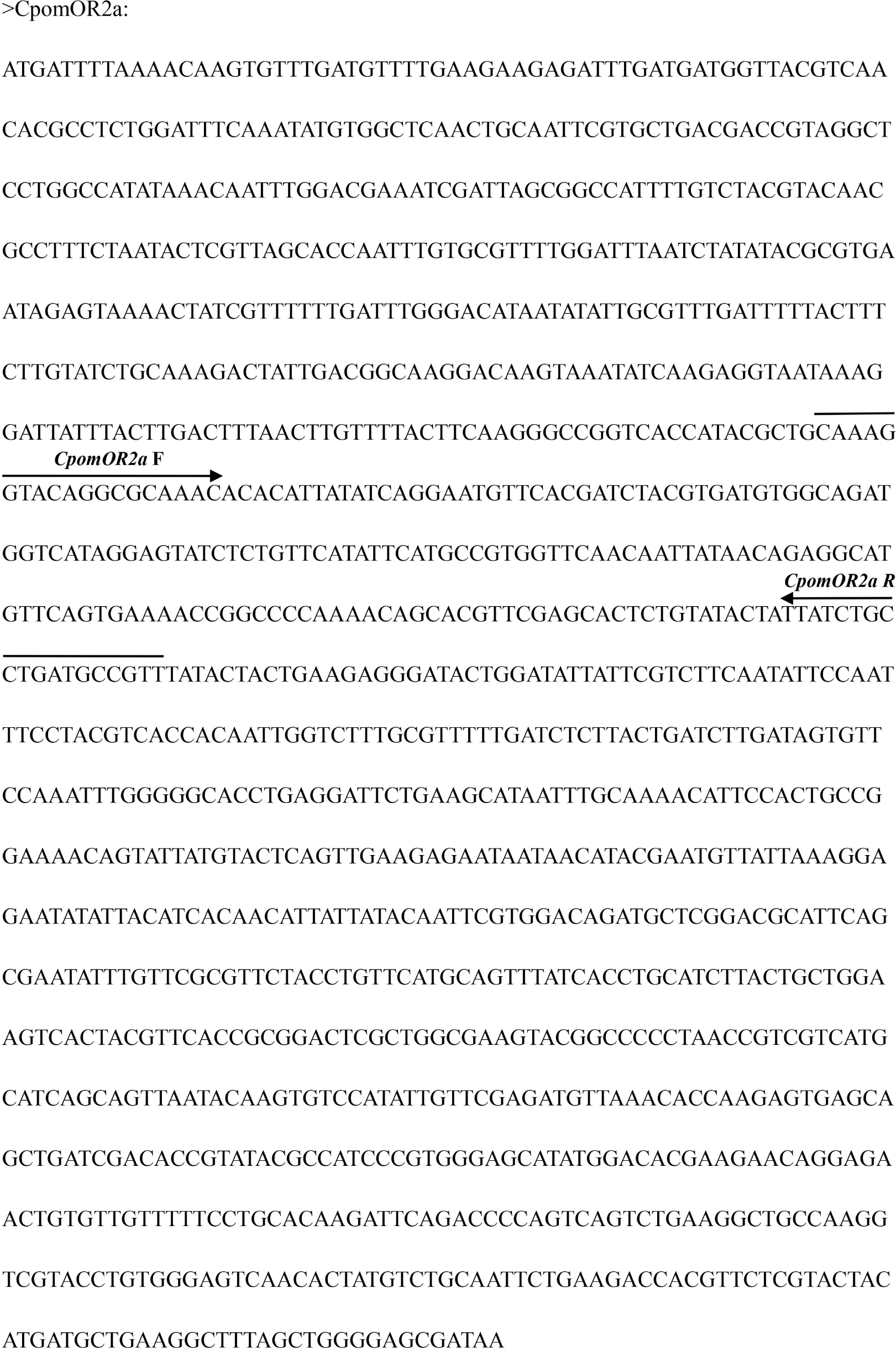
The cDNA sequence of CpomOR2a.The location of the primer sequence is indicated by an arrow.

**Figure S6.**
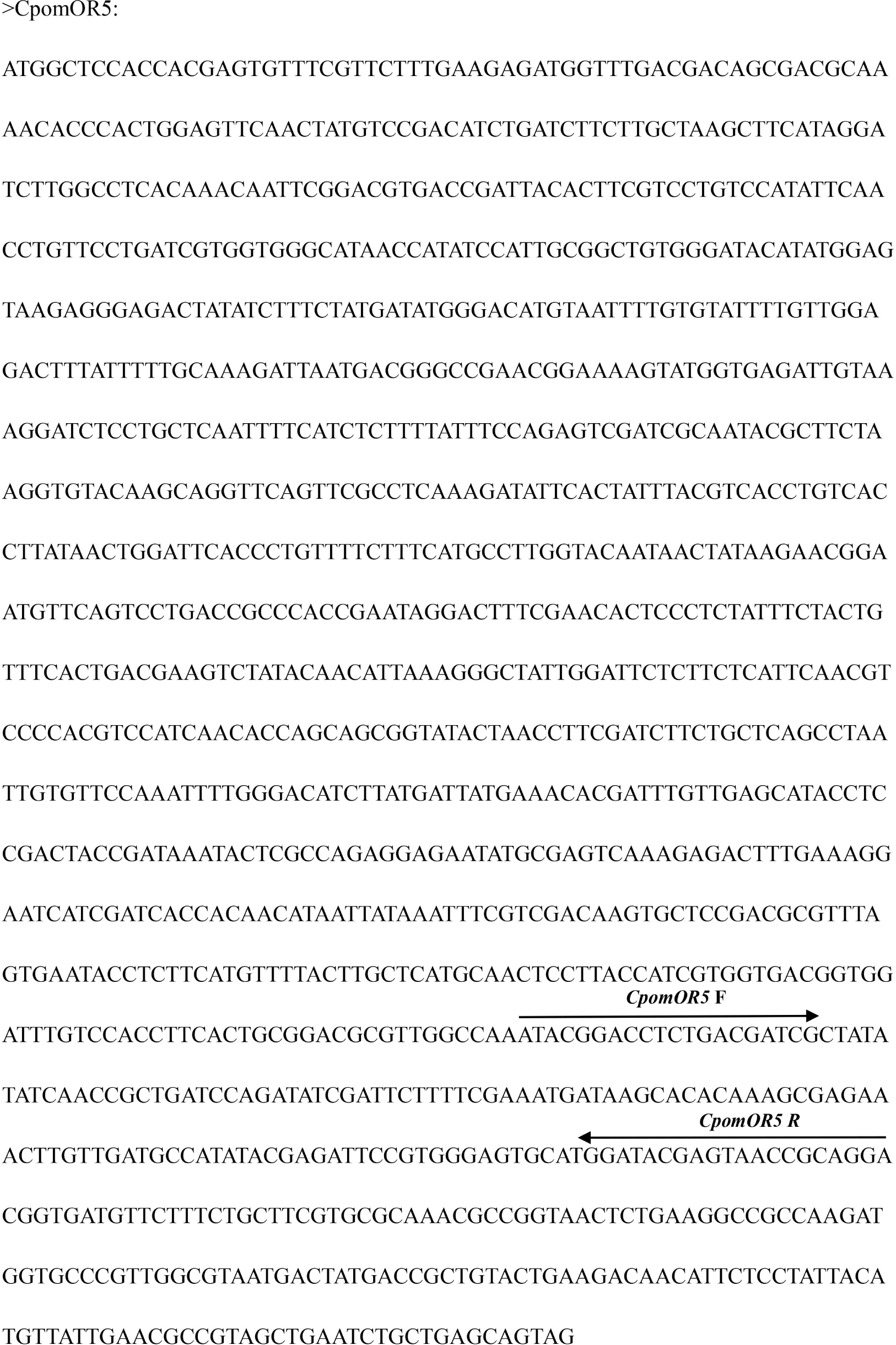
The cDNA sequence of CpomOR5.The location of the primer sequence is indicated by an arrow.

**Figure S7.**
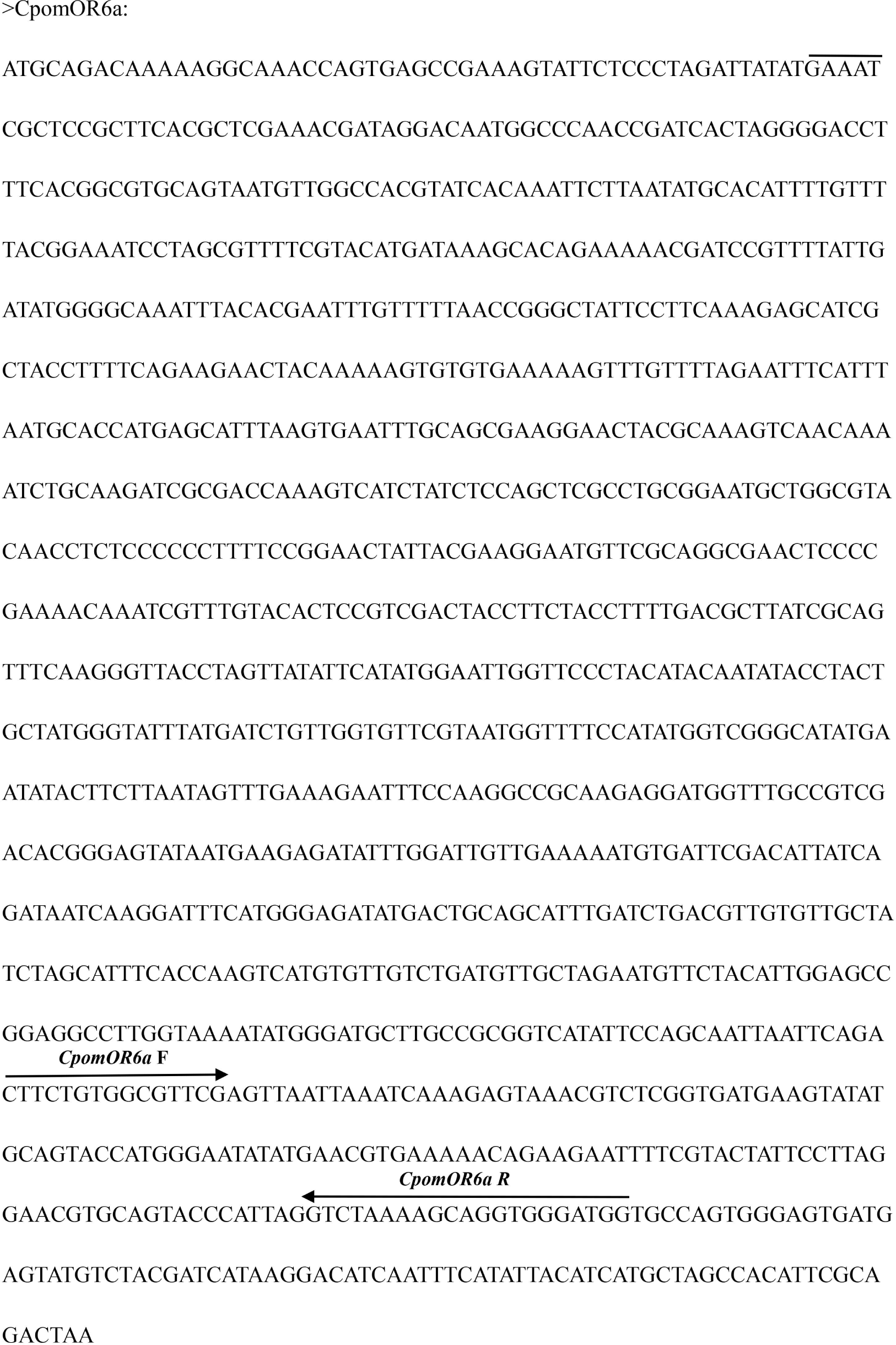
The cDNA sequence of CpomOR6a.The location of the primer sequence is indicated by an arrow.

**Figure S8.**
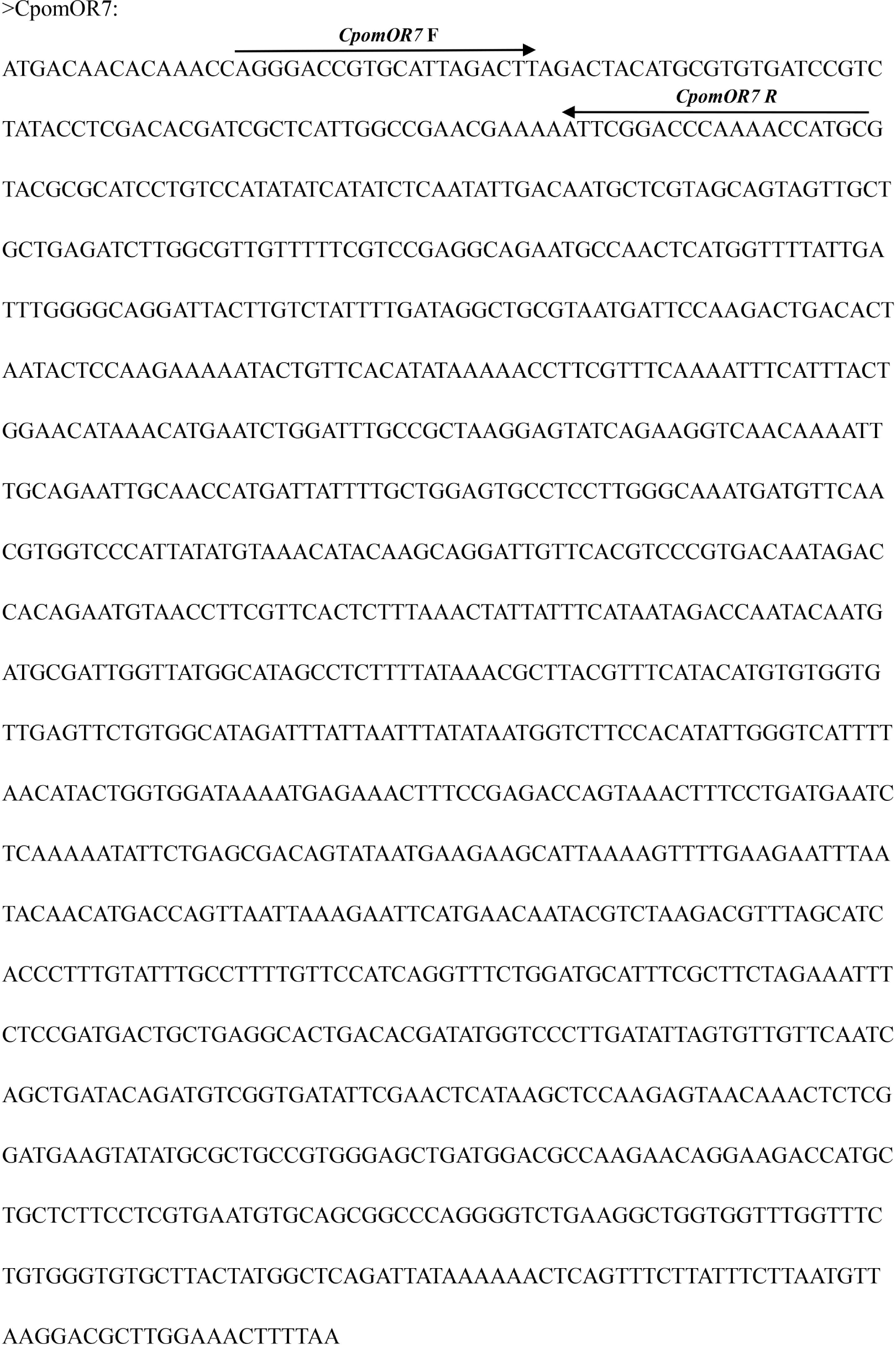
The cDNA sequence of CpomOR7.The location of the primer sequence is indicated by an arrow.

**Figure S9.**
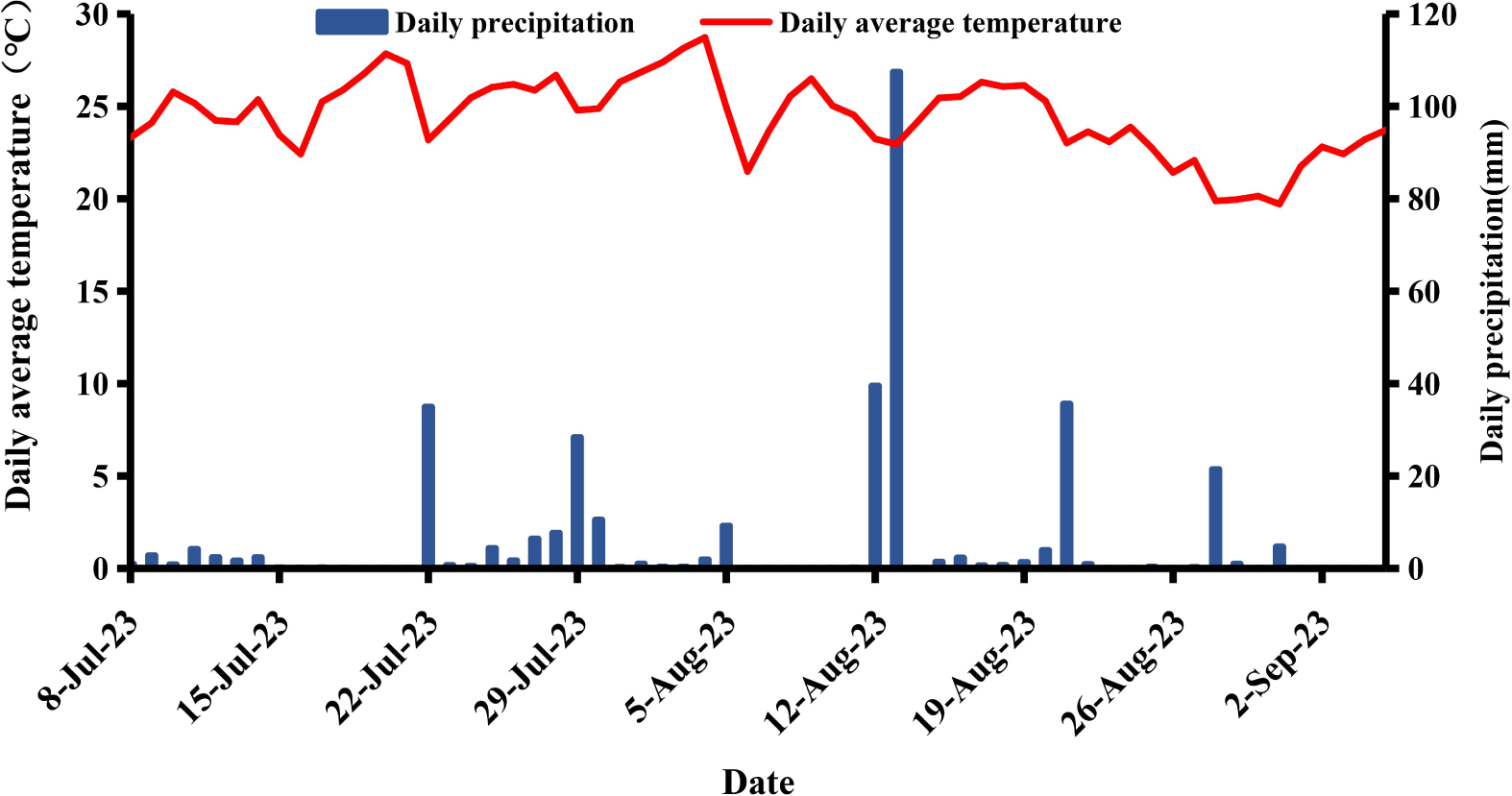
Thee ambient temperature and precipitation event at the experimental site throughout the experiment.

**Figure S10.**
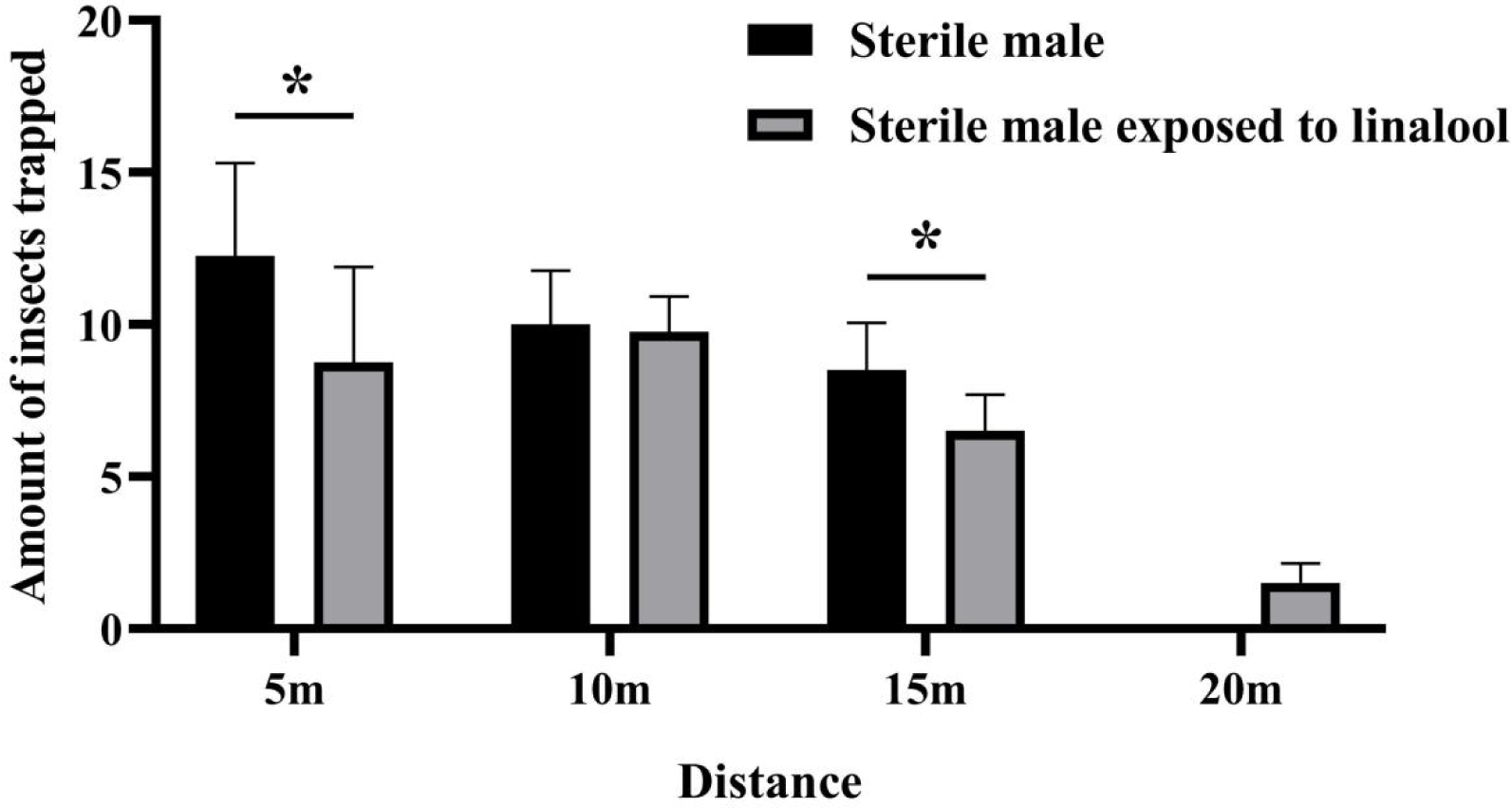
The number of sterile males and fumigation-treated sterile males of *C. pomonella* recaptured at different distance (m) far away from the release center. Asterisk (*) on the error bars indicate significant differences analyzed by the one-way analysis of variance (ANOVA) with independent samples t-tests (*, P < 0.05).

